# Assessing bivalve phylogeny using Deep Learning and Computer Vision approaches

**DOI:** 10.1101/2021.04.08.438943

**Authors:** Steffen Kiel

## Abstract

Phylogenetic analyses using morphological data currently require hand-crafted character matrices, limiting the number of taxa that can be included. Here I explore how Deep Learning and Computer Vision approaches typically applied to image classification tasks, may be used to infer phylogenetic relationships among bivalves. A convolutional neural network (CNN) was trained on thousands of images showing species of 75 bivalve families. The predictions of the CNN on a large number of bivalve images are then interpreted as an indication of how similar these bivalves are to each other, are averaged by the families to which the species belonged, and visualized in a cluster diagram. In this cluster diagram, significantly more families clustered with members of their subclasses than expected by chance, confirming the feasibility of the approach. To address the issue of convergent evolution, two further CNNs were trained, on the same images but grouped by the orders and subclasses to which the species belonged. Combining predictions for the same images but on different taxonomic levels improved the inferred phylogenetic relationships also of families that the CNNs had not been trained on. Finally, this combined tree is merged with five published phylogenetic trees into a supertree, representing the largest single phylogeny of the Bivalvia to date, encompassing 128 families, including six exclusively fossil families and nine extant families for which presently no molecular data are available. Issues inherent to the approach and suggestions for future directions are discussed.

## INTRODUCTION

The fossil record is the primary archive of life’s history from local to global scales, providing key evidence for rates and patterns of evolution (Foote et al. 2007; Jablonski et al. 1983; Knope et al. 2020) and the history of biodiversity through geological time (Peters & Foote 2001; Renema et al. 2008; Sepkoski 1981). A major model group for such evolutionary studies are bivalve mollusks, due to their ubiquity in modern and ancient oceans (Bieler et al. 2014), the strong link between form and function of their shells (Stanley 1970), and the fidelity of their fossil record (Jablonski & Finarelli 2009; Valentine et al. 2006). They have for example been used in studies of large-scale faunal replacements through the Phanerozoic (Gould & Calloway 1980), patterns of survival across major mass extinction events (Lockwood 2003; Raup & Jablonski 1993), and the evolutionary dynamics of global latitudinal diversity gradients (Crame 2002; Jablonski et al. 2006).

However, much of this work used quantitative methods and little attention was paid to phylogenetic history or organism relationships (Patzkowsky 2017). While the molecular revolution has dramatically improved our understanding of the phylogenetic relationships among and within the major extant animal clades (Dunn et al. 2008; Laumer et al. 2019; Pick et al. 2010), about 99% of all animals that ever inhabited Earth are extinct (Erwin 2008) and hence inaccessible to molecular phylogenetic methods. Furthermore, performing a phylogenetic analysis on fossil organisms is labor-intensive as it presently requires a hand-crafted character matrix, and even the largest analyses of entire animal classes rarely use more than 100 taxa (Bieler et al. 2014; Kroh & Smith 2010; Thuy & Stöhr 2016).

Here I explore how computerized image classification may be used to infer phylogenetic relationships. Recent advances in Deep Learning have dramatically improved the state-of-the-art in visual object recognition and image classification, and these methods are now used in a wide range of fields including face recognition, cancer diagnosis, self-driving cars, and many others (LeCun et al. 2015). In the geological and paleontological sciences these approaches are used in seismics (Geng et al. 2020; Huang et al. 2017a; Huang et al. 2017b), lithofacies classification (Baraboshkin et al. 2020; de Lima et al. 2020a; de Lima et al. 2019; John & Kanagandran 2019; Koeshidayatullah et al. 2020), and the identification of index fossils (de Lima et al. 2020b). Applications in the biological sciences focus on automated taxon identification (Hsiang et al. 2019; Mata-Montero & Carranza-Rojas 2016; Wäldchen & Mäder 2018; Wäldchen et al. 2018) where accuracies for certain plant and insect groups reach up to 99.9% (Barré et al. 2017; Valan et al. 2019). Images of organisms, including their fossils, are becoming increasingly available through large-scale digitization efforts of museum collections like iDigBio (https://www.idigbio.org/) and DiSSCo (https://www.dissco.eu/), and are made accessible through bioinformatics infrastructures such as GBIF (https://www.gbif.org/). Furthermore, the means to automatically resolve the validity of taxonomic names are rapidly increasing thanks to global, authorative databases such as the World Register of Marine Species (WoRMS: http://www.marinespecies.org/index.php).

## THE CONCEPT

In a typical image classification task, a convolutional neural network (CNN) learns to classify images belonging to predefined categories. To achieve this, the CNN analyzes a large set of images belonging to these categories, to find general features that the images of each category have in common. For the actual classification of a given test image, the trained CNN returns a set of predictions (probabilities) by which the image belongs to each of the categories the CNN was trained on: the categories with the highest values are those that ‘look similar’ to the test image, and with decreasing prediction values, the categories will look increasingly different. When applying this concept to images showing the morphology of bivalve shells, the prediction values should give an indication how morphologically similar the bivalve on the test image is to other bivalves. Thus, when predictions are obtained for images encompassing a wide range of bivalve clades, these predictions can be regarded as a distance matrix of the morphological similarities of the bivalve clades in question. Based on the – certainly naïve – assumption that morphologically similar taxa are phylogenetically related, visualizing the distance matrix as a cluster diagram could provide insights into the phylogenetic relationships among the taxa.

Any evolutionary biologist or taxonomist reading these lines will immediately think of convergent evolution – the independent evolution of similar morphological features among different clades. Two approaches are explored here to address this issue, both taking advantage of existing phylogenetic information. The *ensemble* approach combines predictions made by several CNNs trained on different taxonomic levels. This approach assumes that features learned to predict higher taxonomic categories also improve the family-level clustering of the families the CNN was not trained on. In the *supertree* approach, the tree derived from the predictions of the CNNs is combined with published phylogenetic trees into a supertree (Ragan 1992).

## MATERIALS AND METHODS

### Data collection and cleaning

The images used here were accessed through three groups of sources: data aggregation platforms such as GBIF and iDigBio, natural history museums, and websites of shell dealers and private enthusiasts (see Table 1 for full list and websites). Also included were images from ongoing digitization efforts of the collection in the author’s care (Swedish Museum of Natural History, Dept. of Palaeobiology). These images commonly showed more than one specimen, or more than one view of a specimen, and often contained other items such as trays, boxes, labels, logos, and a diversity of scale bars, including coins. To maximize the information density of the images, and to reduce noise and potential bias caused by objects other than bivalves, all images were subject to an automated image segmentation process to decompose them into their individual items. The segmentation was done with a thresholding approach (Sahoo et al. 1988) using the R packages imager (Barthelmé & Tschumperlé 2019) and magick (Ooms & Team 2020). Only images showing the inner or outer lateral side of the shells were kept. When necessary, images were rotated into the correct scientific position with the hinge line up, to the best possible extent by steps of 90°.

**Table 1.**
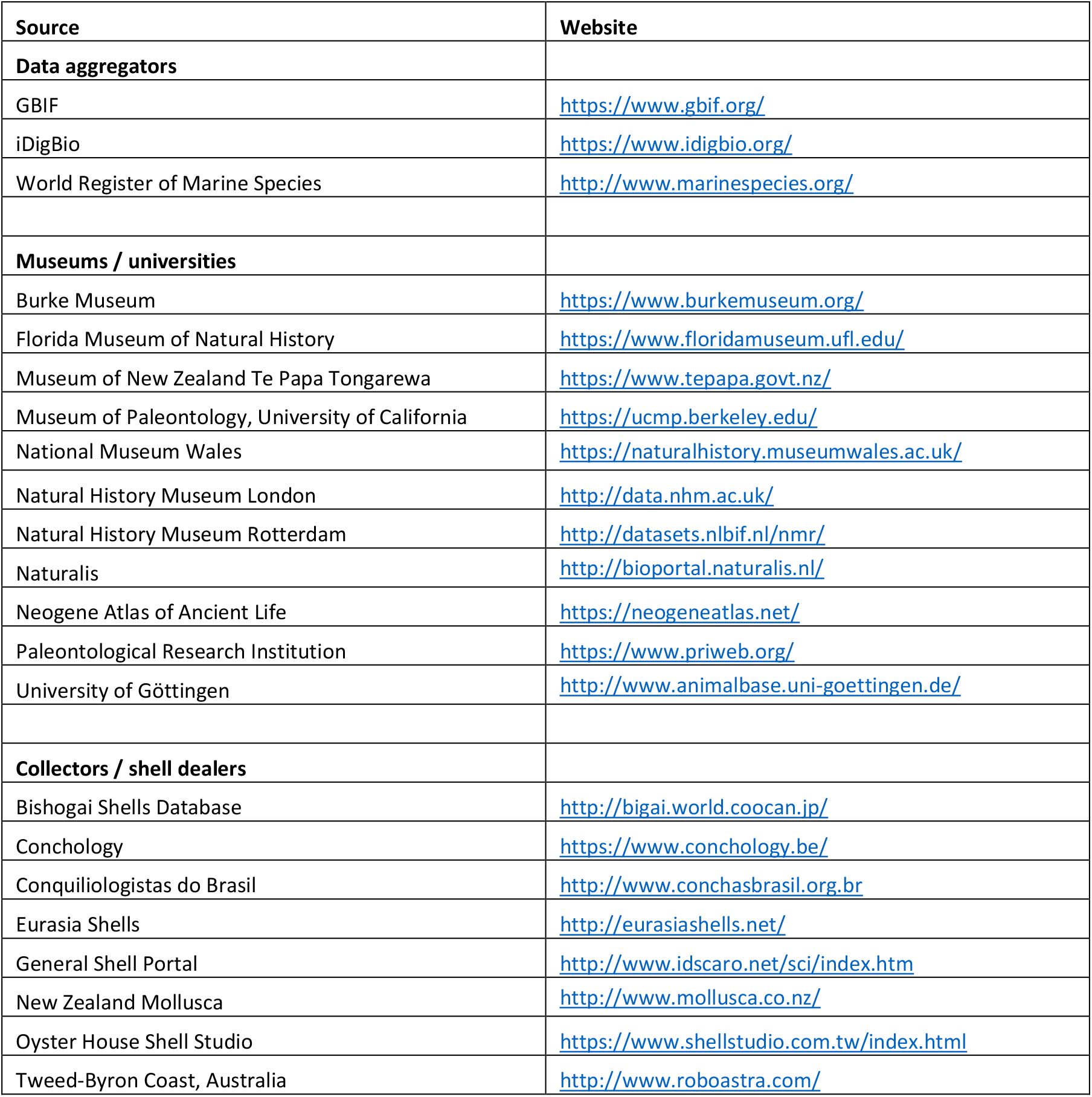
Image sources.

Taxonomic control was as follows: for extant taxa, only images with available species-level identification were used. The accuracy of the identification on family level was first assessed based on my own taxonomic expertise, and any image that I considered incorrectly identified on the family level was removed. Then each species-level identification was checked against WoRMS (as of September 2020): when necessary, the species were re-assigned to the species/genus/family to which they belonged according to WoRMS; images of species not found in WoRMS were removed. Although this cleaning removed a sizeable number of images from the training data set, it improved the prediction accuracy of the CNNs. For images of fossil taxa, an identification to family level had to suffice due to the still low coverage of fossil species in WoRMS. The only taxonomic check was my own evaluation on the level of family. The classification on the level of order followed WoRMS, and if no order was available, the superfamily indicated in WoRMS was used. The assignment to a subclass did not follow WoRMS, instead, the more traditional classification into Protobranchia, Pteriomorphia, Palaeoheterodonta, Archiheterodonta, Anomalodesmata, and Imparidentia, used in many recent phylogenetic publications on the Bivalvia was used (Bieler et al. 2014; Combosch et al. 2017).

The final image dataset included 74820 images belonging to 114 families. All families, their classification, image numbers, use for training, and presence in the phylogenies for the supertree, are listed in the supplemental Table S1. The dataset is highly imbalanced, with a few families being represented by several throusand images, and about half of the families by less than 200 images (Figure 1).

**Figure 1.**
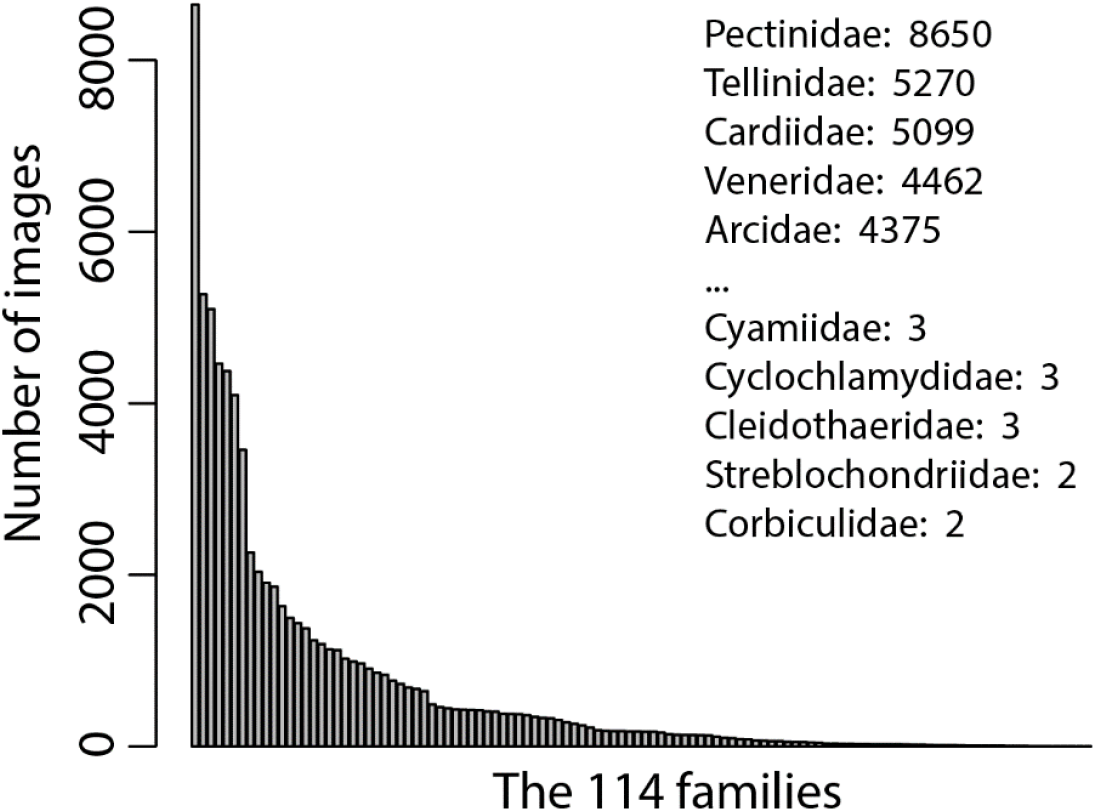
Frequency distribution of bivalve families in the image data set; also shown are image numbers for the five most abundant and five rarest families.

### Training the CNNs

Convolutional neural networks used in Deep Learning consist of numerous layers stacked upon one another. A layer consists of numerous neurons (hence the name neural network), each of which performs some transformation of the input data, for example an image. The result of this transformation is given a certain weight based on how important the output of this neuron was in correctly predicting the category of the input data, and is then passed on to the next layer. This way each successive layer learns increasingly complex features of the data. The final layer of the CNN outputs a prediction to which of the categories the input data (here, the image) belongs. This prediction is evaluated by a ‘loss function’ and depending on whether the prediction was right or wrong, the loss function updates the weights of each neuron in the CNN, a method that is called ‘backpropagation’.

The entire process is iterated numerous times on all available data, and each iteration is called an ‘epoch’. To monitor the learning process, predictions on a different set of data (called validation data/images) are made at the end of each training epoch. The end of the training process is reached when the accuracy of the predictions on the validation data stops to increase compared to previous epochs. At this point, the CNN can be put to use on actual classification tasks.

When trained on millions of images of a wide range of objects, most layers of a CNN learn very general features of these objects, and that such a ‘pre-trained’ CNN can be adopted to solve a more specialized problem. This technique is called ‘transfer learning’: a pre-trained CNN is use as basis and only its last layer, or last few layers, are trained on the specific task at hand (Azizpour et al. 2016; Bengio 2011; Yosinski et al. 2014). This does not only reduce the training time, it also makes it possible to train on much smaller sets of images, because the pre-trained CNN has already learned general features such as edges, line, curves, corners, and other basic shapes. For image classification purposes, various CNNs trained on ImageNet (Deng et al. 2009) on millions of images are available, including VGG16 (Simonyan & Zisserman 2014), Xception (Chollet 2016), and Inception (Szegedy et al. 2016).

Here the ResNet50 V2 architecture (He et al. 2015) is used. Only the final layer of the ResNet50 CNN is trained, using the stochastic gradient descent optimizer (Sutskever et al. 2013); the chosen hyperparameters are shown in Table 2. All images were re-scaled to 224*224 pixels, which is the standart input size for ResNet50. To increase the number of training images during the training, data augmentation was used (Perez & Wang 2017): copies of each image were subject to one or several random transformations, including rotation to up to 30 degrees, shifts in the color channels, a zoom into the image of up to 20%, horizontal and vertical shifts up to 20% of image size, shearing of up to 10 degrees, and horizontal flip. To counter the effect of the highly imbalanced distribution of images among the training categories (Figure 1), the updates during backpropagation were weighted in inverse proportion to the total number of images of the respective category. In other words, the updates of categories with only few images were given higher weight than those of categories with abundant images.

**Table 2.** Setup of the three CNNs and hyperparameters of the stochastic gradient descent optimizer.

| Taxonomic level | Categories in final layer | Learning rate | Decay | Momentum |
| --- | --- | --- | --- | --- |
| family | 75 | 0.01 | 0.001 | 0.9 |
| order | 25 | 0.1 | 0.01 | 0 |
| subclass | 6 | 0.01 | 0.001 | 0.5 |

Another measure to mediate the effect imbalanced distribution of images was to use only families with 60 or more images. This limited the number of families used for training to 75 and these families are henceforth referred to as ‘training families’. The total number of images of the training families was 74069. The remaining 39 families with fewer than 60 images (total 751images) were used only for predictions and henceforth called ‘target families’. The position of these target families on the ensemble and supertrees are of particular interest, because these are the ones that the present approach places on the trees without prior knowledge of their phylogenetic affinities. In case of the supertree, the families present in any of the publish phylogenies used to construct the supertree were excluded from the target families, so that the supertree included 21 target families.

Three individual CNNs with almost identical architecture were each trained on the same 74069 images of the training families, but grouped at different taxonomic levels. The difference between the three CNNs was in the number of prediction provided by the final output layer: the family-CNN provided predictions for 75 families, the order-CNN for 25 orders, and subclasses-CNN for six subclasses. Each CNN was trained for 600 epochs and the weights of each epoch were saved for later selection of the most suitable training epoch (see next section).

### Constructing and selecting trees

The phylogenetic trees were constructed as follows:

1. Predictions were obtained for all available images (both training and target),
2. the predictions were averaged by the families to which the images belong to, so that one averaged prediction value was available for each of the families,
3. these averaged prediction values were used to calculate a distance matrix based on the Bray-Curtis distance measure,
4. the distance matrix was then hierarchically clustered using the complete linkage method.

This approach has an important implication for the selection of the best training epoch of the CNN: The learning success of the CNN after each training epoch is measured by how well it predicts the categories of the evaluation images (see section *The Concept* above). A prediction is considered correct when the category with the highest probability matches the actual category of the respective image. However, the clustering of the categories in the distance matrix is essentially based on all *but* the one with the highest probability. Indeed, if the CNN would always give the correct prediction, and with 100% probability, the resulting cluster diagram would be a simple rake without any subclusters. A consequence is that the training epoch with the highest validation accuracy is not necessarily the one resulting in the phylogenetically most coherent clustering.

To address this issue, the saved weights of each of the 600 training epochs were successively re-loaded into the CNN and predictions on a reduced (to save computation time) set of images were obtained for each epoch. These predictions were then used to reconstruct a tree for each training epoch, and a set of evaluation indices (described below) was calculated for each tree. The epochs were ranked by each of these indices, and based on the lowest sum of ranks, the best overall training epoch was identified. Finally, predictions on the full set of images were obtained using the weights of the best training epoch, and were used to produce the final tree. The evaluation indices are:

- Tree coherency index. A desired property of the tree is that as many tips as possible are included in a few monophyletic subtrees (i.e., that the major clades are retrieved as monophyletic). ‘Monophyletic’ for the present purpose means belonging to the same subclass. Monophyletic subtrees nested within larger monophyletic subtrees were not included in the count. The index was calculated as follows: (number of tips in non-nested, monophyletic subtrees / number of non-nested, monophyletic subtrees) * (number of tips in non-nested, monophyletic subtrees / total number of tips).
- Node distance from tip to nearest kin. The ‘node distance’-indices give the average number of nodes between the tips (families) and the nearest member of the order or subclass they belong to; as ‘members’ qualified only the training families but not the target families. The indices are calculated separately for training and target families.

An issue with the tree coherency index is that it could give quite misleading values due to just a few misplaced families: a single misplaced family in a large and otherwise monophyletic subtree has the potential to drastically reduce the number of tips counted as belonging to monophyletic subtrees (Figure 2A). A disadvantage of the node distance index is that it would suggest good consistency even for rake-like trees composed of pairs each made of two families belonging to the same clade (Figure 2B). Hence, for the selection of the ‘best’ tree, the node distances of the training families and the tree coherency index are here considered in concert. The node distance indices for the target families were only used for subsequent assessments of how close they were placed to members of the order/subclass to which they are currently assigned according to WoRMS.

**Figure 2.**
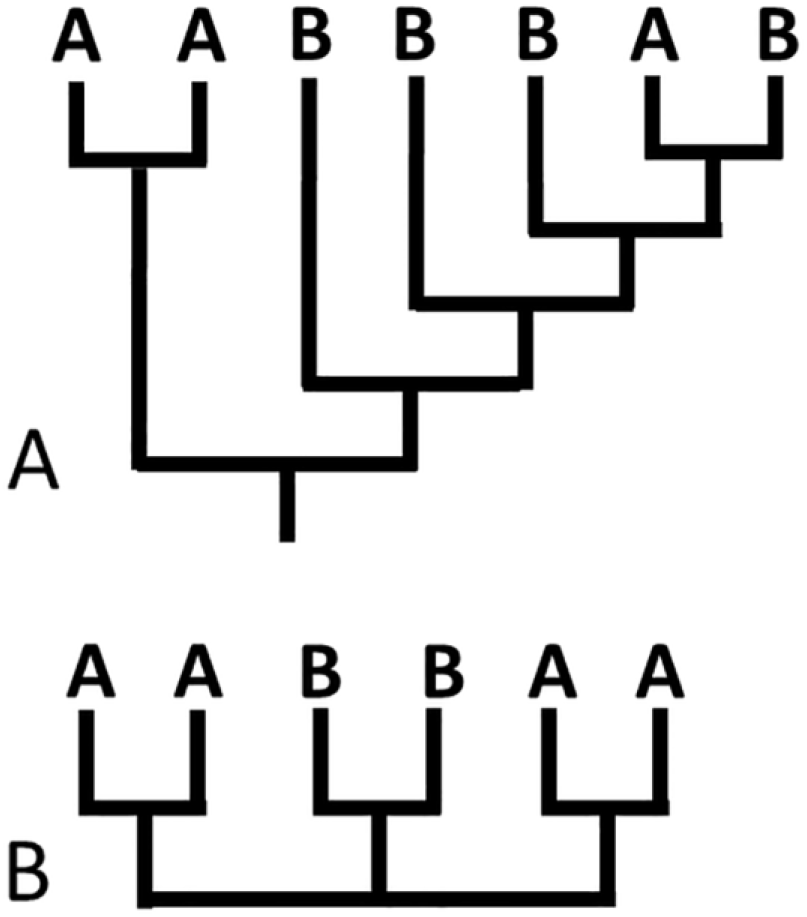
Issues with the indices for tree consistency. **A.** Due to a single misplaced tip, the tree coherency index would rank this tree poorly because only two out of seven tips belong to a monophyletic subtree. **B.** The node distance index would give this tree the highest possible ranking because each tip is sister to one of its kin.

### Constructing the ensemble tree

To construct the ensemble tree, predictions were obtained from each of the three CNNs for all available images (both training and target families). For each of the three sets of predictions, the predictions were averaged by the families to which the images belonged. Thus the set of averaged predictions of the family-CNN included for each family the probabilities by which it belonged to the each of the families the family-CNN was trained on; the predictions set of the order-CNN included for each family the probabilities by which it belonged to the each of the orders the order-CNN was trained on, and so forth. To combine the three sets of averaged predictions, the averaged predictions of the family-CNN were used as base, and the following steps were carried out for each prediction:

1. The order to which the predicted family belongs was identified (Figure 3A),
2. the prediction for that order was looked up in the predictions set of the order-CNN (Figure 3A),
3. that value was multiplied by a given factor (see below) and then added to the prediction of each training family belonging to that order (Figure 3B).
4. The same updates were done for the subclasses.

**Figure 3.**
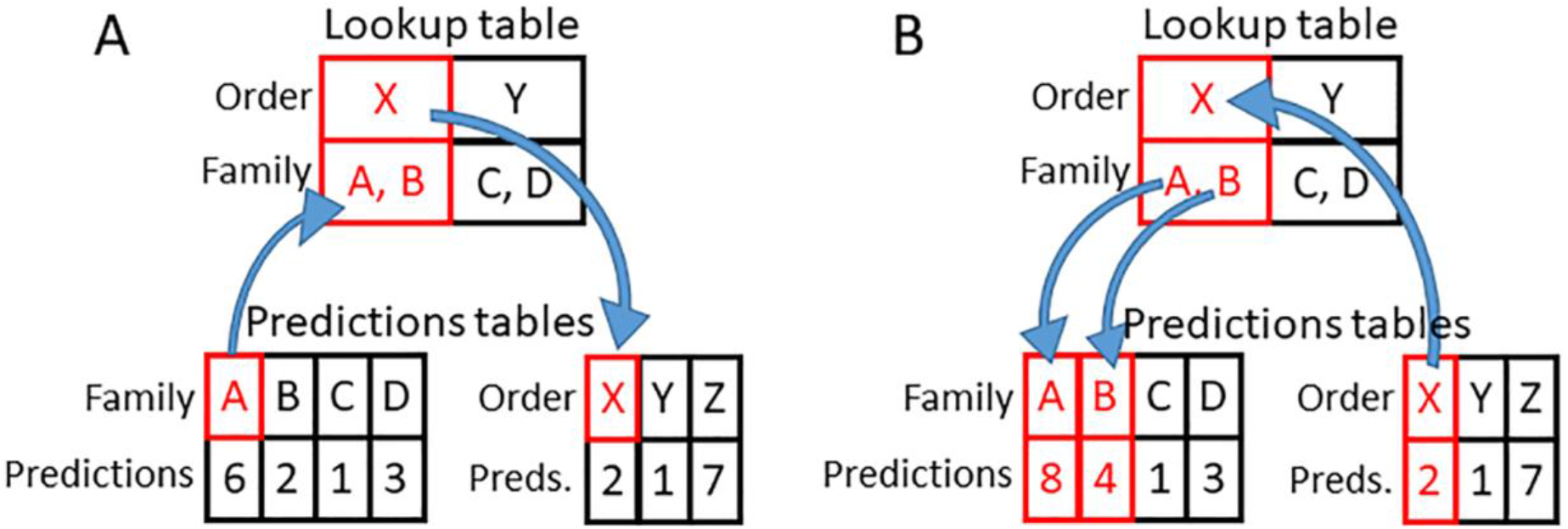
Schematic illustration how predictions from different taxonomic levels were combined for the ensemble tree. **A.** Identifying the order the current family belongs to. **B.** Adding the prediction value for that order to the prediction value of each of its families.

Combining the three sets of predictions in equal proportions does not necessarily result in the most accurate tree. For example, features learned to distinguish subclasses could potentially distort or mask more fine-grained distinctions on the family level. Therefore, a series of different proportions was tested, by multiplying each order- and subclass-level prediction with a given factor (as described above in step 3) before adding it to the family-level prediction. For each combined predictions set in this series, a tree was constructed and assessed using the tree coherency index and node distance indices for the training families, described in the *Constructing and selecting trees* section above. Then the trees were ranked by each of these indices, and based on the lowest sum of ranks, the best overall tree was identified.

### Constructing the supertree

For the supertree, five phylogenies that covered the entire class Bivalvia were used, four of which were based exclusively on molecular data (Combosch et al. 2017; Gonzáles et al. 2015; Plazzi & Passamonti 2010; Sharma et al. 2012), and one included also conchological, morphological, and anatomical characters (Bieler et al. 2014). The published trees were reconstructed semi-manually using TreeSnatcher plus (Laubach et al. 2012), and were combined with the ensemble tree using matrix representation (Ragan 1992).

### Code

All code related to the CNNs (training, predictions, Grad-CAM) was written in Python using Keras (Chollet 2015) with a TensorFlow backend (Abadi et al. 2016). All phylogenetic reconstructions and tree analyses were performed in R using the packages adephylo (Jombart et al. 2010), ape (Paradis et al. 2004), ctc (Lucas & Gautier 2020), dendextend (Galili 2015), phangorn (Schliep 2011), and vegan (Oksanen et al. 2005). The code is available on github (https://github.com/steffenkiel/bivalve_phylo).

## RESULTS & DISCUSSION

### Tree topology through training

The best overall tree by the family-CNN was found early during the training, after about one fifth (127^th^ epoch; Table 3) of the training. Noteworthy in the development of the tree evaluation indices through training is that the node distances of the training families do not improve much through the training period, but those of the target families do (Figure 4A). The trees of the order-CNN show the reverse trend: the node distances of the training families to their respective orders and subclasses improved notably, while those of the target families did not; they improved only during the first 50-70 training epochs and dwindles afterward (Figure 4B; Table 3). Among the trees of the subclass-CNN, the node distances of the training families to their subclasses improved through training, the best values for their orders were reached during the first 100 training epoch and then remained almost constant at a slightly worse level for the rest of the training. For the target families, the node distances to their orders improved slightly through training, while they remained nearly constant for the subclasses (Figure 4C; Table 3).

**Table 3.** Summary of training results: Prediction accuracies and tree consistency indices for 600 epochs, for each of the three CNNs.

| Taxonomic level | Epoch with highest prediction accuracy | Highest prediction accuracy | Epoch with highest TCI | Epoch with best ND to order | Epoch with best ND to subclass | Best epoch overall |
| --- | --- | --- | --- | --- | --- | --- |
| family | 476 | 0.69 | 514 | 3 | 188 | 127 |
| order | 396 | 0.64 | 95 | 499 | 331 | 530 |
| subclass | 344 | 0.77 | 433 | 51 | 222 | 483 |

**Figure 4.**
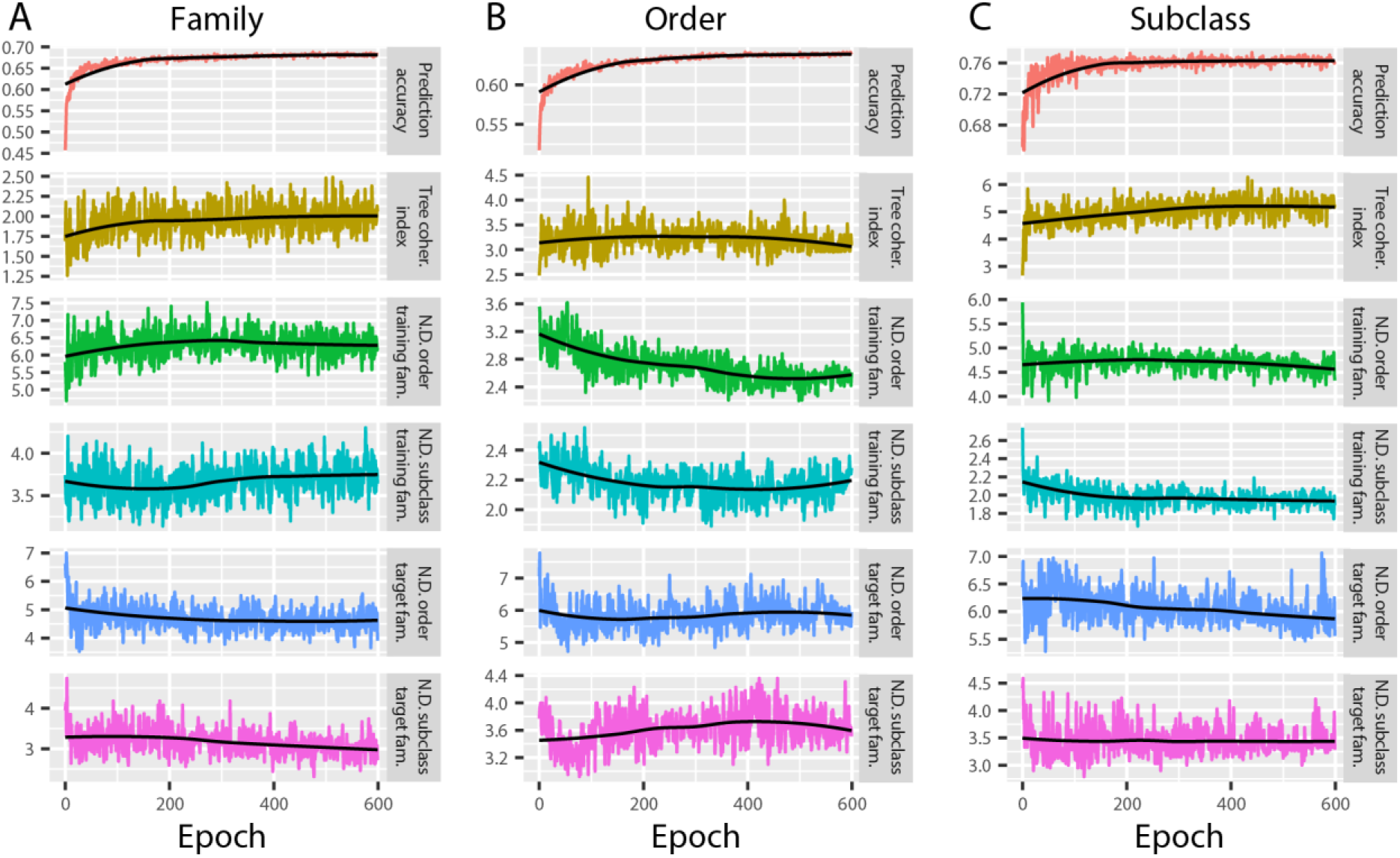
Development of validation accuracy and tree consistency indices during 600 epochs of training, for the CNNs trained on family (**A**), order (**B**), and subclass (**C**) level. N.D. = node distance.

The family-CNN shows a lack of improvement of the node distances among training families during training (Figure 4A). This might indicate that the limits of what the shell morphology on the available images can tell about order and subclass memberships of bivalve families have been reached. This does not necessarily indicate, however, that also the limits to what shell morphology can tell about order and subclass memberships of bivalve families has been reached.

During training of the order-CNN and subclass-CNN, the node distances of the training families to their respective orders and subclasses improves notably (Figure 4B, C). This is expected because the more accurate the order- and subclass-CNNs become in classifying images into their orders and subclasses, the closer are their respective families to each other in the tree. However, continued training of the subclass-CNN did not improve node distances of the training families to their order, indicating that morphological features learned to recognize subclass membership do not help in assigning the family to the correct order within the subclass. Vice versa, training of the order-CNN improved node distances to the subclasses only for the first third of the training, but diminished afterward. The initial classification of families to the correct order has the side effect that the families are also placed closer to the members of their subclass: namely to those that are in the same order. But in the long run, learning features to recognize order membership does not help to identify relationships among orders.

Interestingly, the node distances of the target families to both order and subclasses improved during training of the family-CNNs. I find this hard to interpret; perhaps the predictions became more focused with continued training and predictions for ‘wrong’ orders and subclass became less frequent. The order- and subclass-CNNs show almost opposing trends. Whereas in the order-CNN, node distances show an overall deterioration, they improved for the subclass-CNN. The deterioration of node distances in the order-CNN might partially be a side effect of the significant improvements of the node distances of the training families. While prediction accuracy of the training images improves, leading to improved clustering of the training families, the prediction accuracies for the images of the target families are left behind and become less distinctive in respect to the orders to which the target families belong. However, there might also be a deeper underlying reason. A target family can only be recognized as a member of its clade when the morphology of the family is within the range of the morphologies in the training set for that clade. By its nature, a subclass includes a broader morphologic diversity than an order. Therefore, a CNN trained on the subclass level may be more likely to correctly assign target families to their subclasses, than a CNN trained on the more restricted morphological range of an order.

### Features used for classification

When classifying organisms, human taxonomist focus on characters with a biological and/or functional meaning (such as hinge dentition, muscle scars, and pallial line in the case of bivalves). In contrast, a CNN essentially uses sophisticated statistics to find similarities among the training images. Visualizing what the predictions of the CNNs are based on can help understanding what the CNNs have actually learned, especially for predictions on different taxonomic levels. It can also aid in detecting errors or biases in the training process, and can potentially point to characteristics of the shells that are typically not considered by human taxonomists (Valan et al. 2019). A visualization technique for this purpose is Gradient-weighted Class Activation Mapping (Grad-CAM), in which the weights of the final output layer of the CNN are used for highlighting the regions in the image that are important for the prediction (Selvaraju et al. 2017). Here a set of 519 images showing a wide range of bivalve shell shapes and features was selected and the Grad-CAM technique was applied to them with each of the three CNNs.

Overall, the three CNNs did not focus on systematically different features. Rather, the focal areas were overlapping if not identical in the vast majority of the cases (Figure 5A, C, F-H). There was a tendency among the focal areas of the order-CNN and subclass-CNN for being less broad and more defined compared to those of the family-CNN (Figure 5A, C, G). Notably different focal areas between the CNNs were often associated with erroneous predictions (Figure 5B). The most common focal areas were:

- sculpture on the anterior and posterior surface (i.e., Anomiidae, Arcidae, Cardiidae, Corbulidae, Mactridae, Noetiidae, Solenidae; Figure 5B),
- shell outline (i.e., Periplomatidae, Spondylidae, Gryphaeidae; Figure 5C),
- hinge area (i.e., Arcidae, Crassatellidae, Laternulidae, Neilonellidae, Nuculanidae; Figure 5D),
- crenulated inner ventral margin (i.e., Arcidae, Glycymerididae, Limopsidae, Donacidae; Figure 5E),
- ears and notches at the dorsal margin (i.e., Pectinidae, Isognomonidae, Margaritidae, Pteriidae; Figure 5F),
- muscle scars, pallial line and sinus (i.e., Lucinidae, Veneridae, Corbulidae; Figure 5G),
- reflecting stripes on the external surfaces, especially of glossy shells (i.e., Astartidae; Crassatellidae, Galeommatidae; Figure 5H).

**Figure 5.**
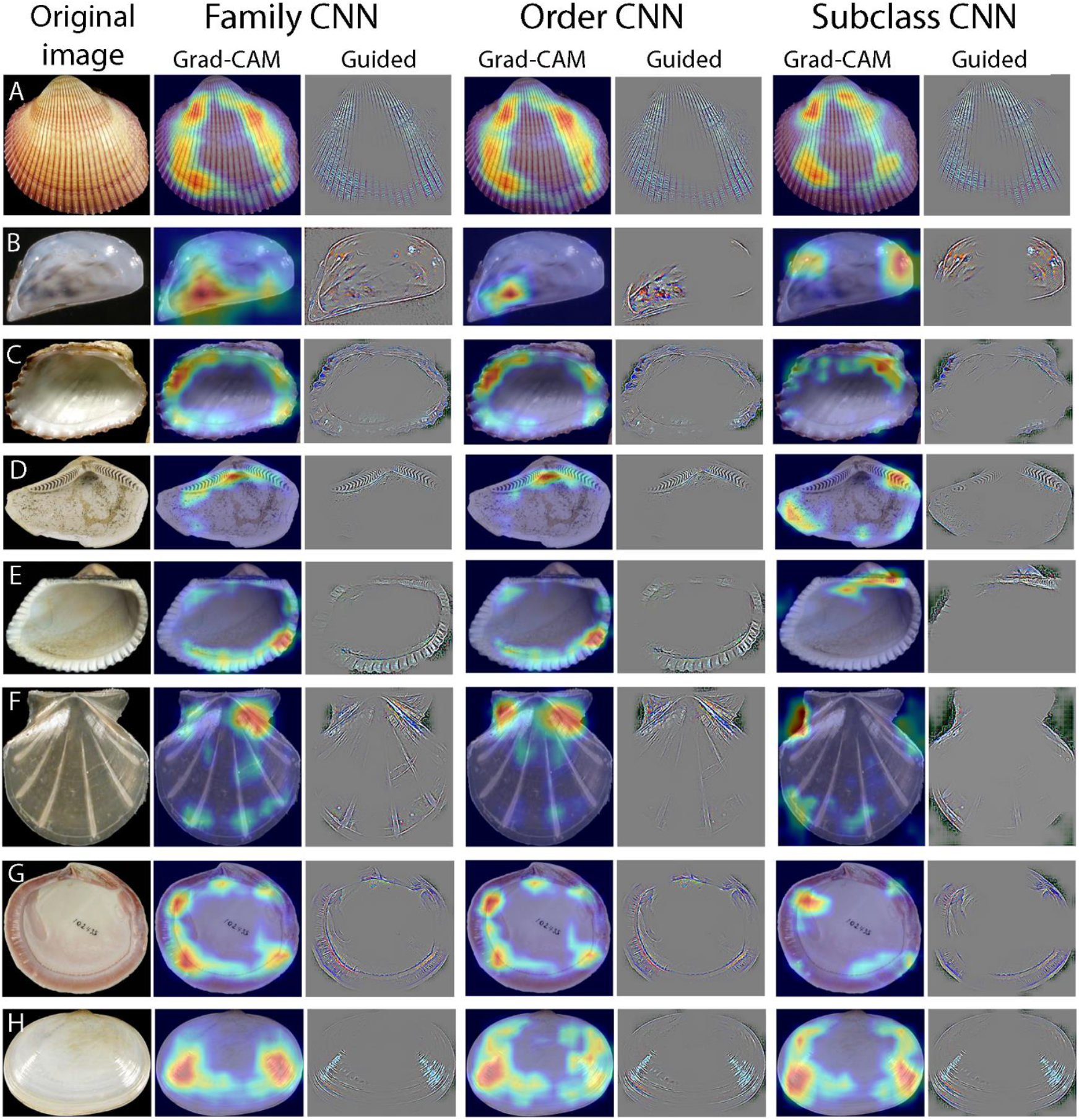
Grad-CAM highlighting the areas on the images most relevant for the CNNs for their predictions. **A**, *Maoricardium pseudolima* (Cardiidae), example for overlapping focal areas between the different CNNs and for focus on ornament near anterior and posterior shell margins. **B**, *Dreissena polymorpha* (Dreissenidae); order misidentified as Mytilida, subclass as Protobranchia). **C**, *Glans afra* (Carditidae), focus on shell outline. **D**, *Nuculana callimene* (Nuculanidae), focus on hinge. **E**, *Anadara kagoshimensis* (Arcidae), focus on crenulated ventral margin. **F**, *Parvamussium ina* (Propeamussiidae), focus on notches at the dorsal margin. **G**, *Codakia distinguenda* (Lucinidae), focus on pallial line and muscle scars. **H**, *Scintilla philippinensis* (Galeommatidae), focus on reflective stripes on outer lateral shell surface. Image sources are www.conchology.be: A, H; University of Göttingen: B; iDigBio: C, F, G; GBIF: D; www.idscaro.net: E.

Note that the lists of families for each point are not exhaustive. Most focus areas are not unlike those used by human taxonomists. Noteworthy are the reflecting stripes on the lateral external shell surfaces. These are perhaps problematic because they are secondary features, resulting from the angle of the light sources used during photography. While it certainly is possible that the curvature of the shell producing the shape and direction of those reflecting stripes is characteristic for a family, these pattern can be different on images taken with light coming from different angles.

### Predictions proportions for the ensemble tree

The best values for the tree evaluation indices of the training families were found when the weight of the contribution of the subclass predictions was 25% or less (Figure 6A, C, E). Whereas the node distance to orders gradually became worse with increasing input from the subclass predictions, the node distance to subclasses remained constant at a good level when the subclass predictions weighted more 25%. In contrast, with increasing weight given to the order predictions, the values of the node distance improved, while the tree coherency index showed a slight decrease (Figure 6B, D, F). The best ensemble tree based on all three indices was found when the contributions of the order predictions weighted 100% and those of the subclass predictions weighted 16%. Thus a small input of subclass predictions appears to ensure that families cluster within their respective subclasses, but too much input does indeed mask the more detailed relationships derived from the family predictions. However, this does not apply to order predictions.

**Figure 6.**
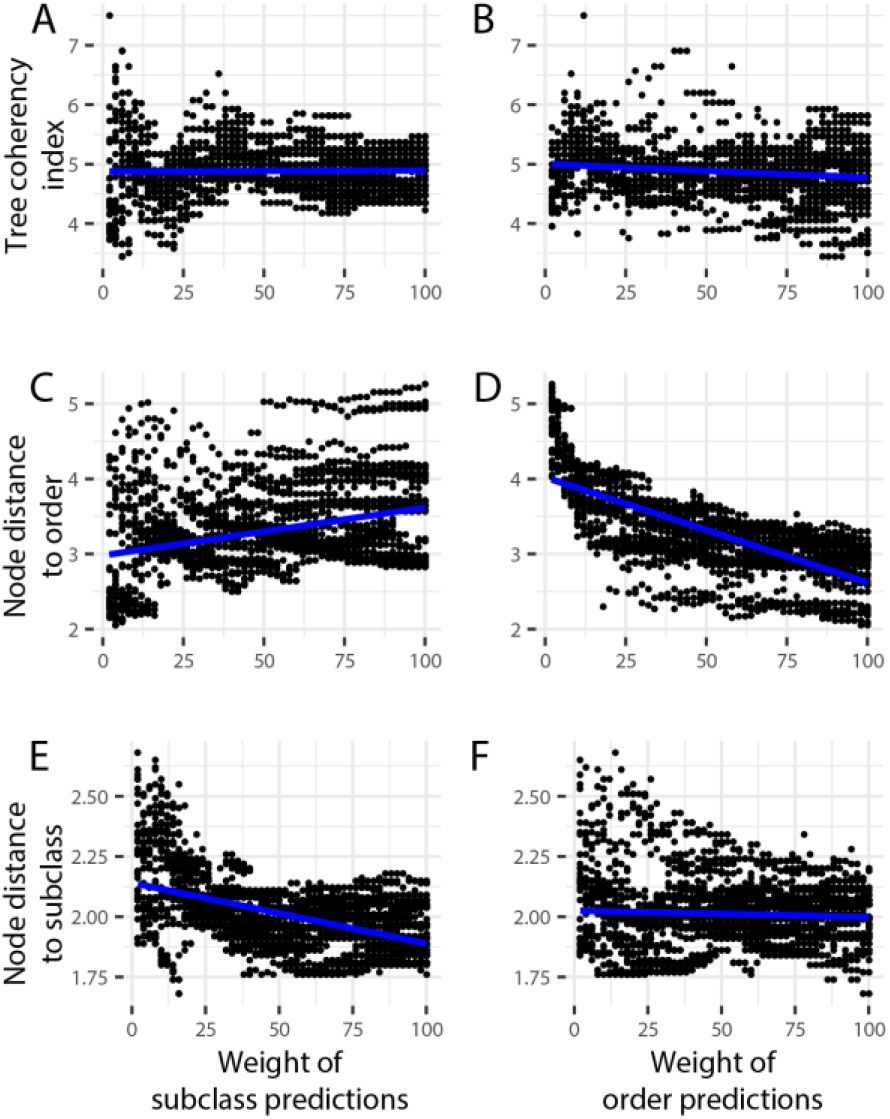
Relationships between the weights of the subclass and order predictions and the tree consistency indices when making the ensemble.

### Phylogenetic relationships and convergence

To facilitate the assessment of the approaches to address the issue of convergent evolution (ensemble and supertree), the tree evaluation indices of three trees were compared: a tree based on predictions of the family-CNN only (including training and target families), the ensemble tree, and the supertree. As a baseline to check if the present approach results in better trees than random clustering, the indices were also calculated for 100 randomized copies of the family-CNN tree (Figure 7).

**Figure 7.**
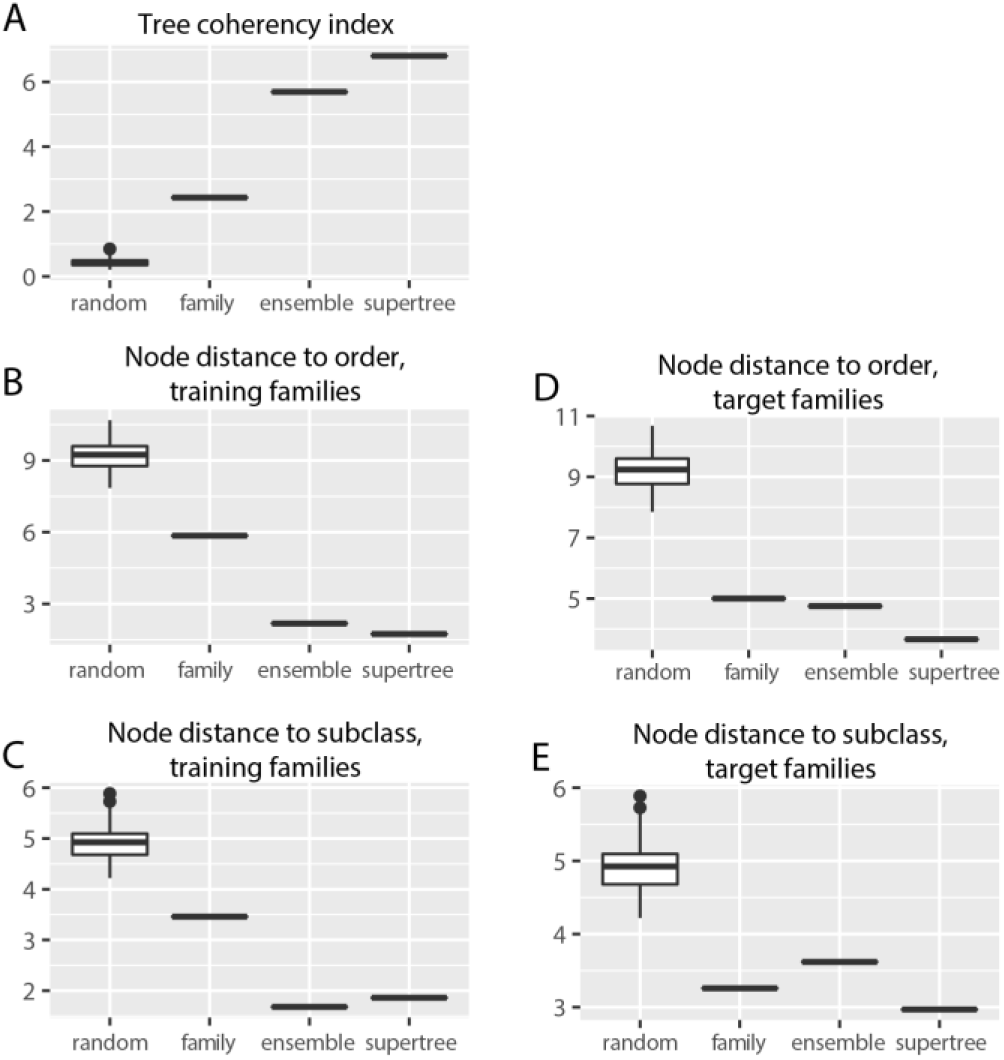
Tree consistency indices of the family, ensemble, and supertree, compared to those of 100 randomized trees of the same size as the family tree.

All trees derived from the predictions of the CNNs reflect phylogenetic relationships better than the best randomized tree. Even the simplest tree derived only from predictions of the family-CNN shows many monophyletic clusters, the largest with ten tips (of Pteriomorphia, including Ostreidae, Pinnidae and Pteriidae), and several with five tips: one of each Protobranchia, Pteriomorphia, and Anomalodesmata, and two of Imparidentia, one of them including Tellinidae and Semelidae, the other including Teredinidae and Xylophagaidae) (Figure 8). These results highlights two things: first, without any prior knowledge of phylogenetic relationships of bivalves, the approach outlined here clusters numerous related bivalve families. Second, however, it also highlights that convergence is indeed a major issue. This is obvious on a large scale because no subclass or order is retrieved as a monophyletic group in the tree derived only from predictions of the family-CNN. It can also be seen on smaller scales, nicely illustrated by the clustering of the very elongated families Solemyidae (Protobranchia) and Solenidae (Imparidentia).

**Figure 8.**
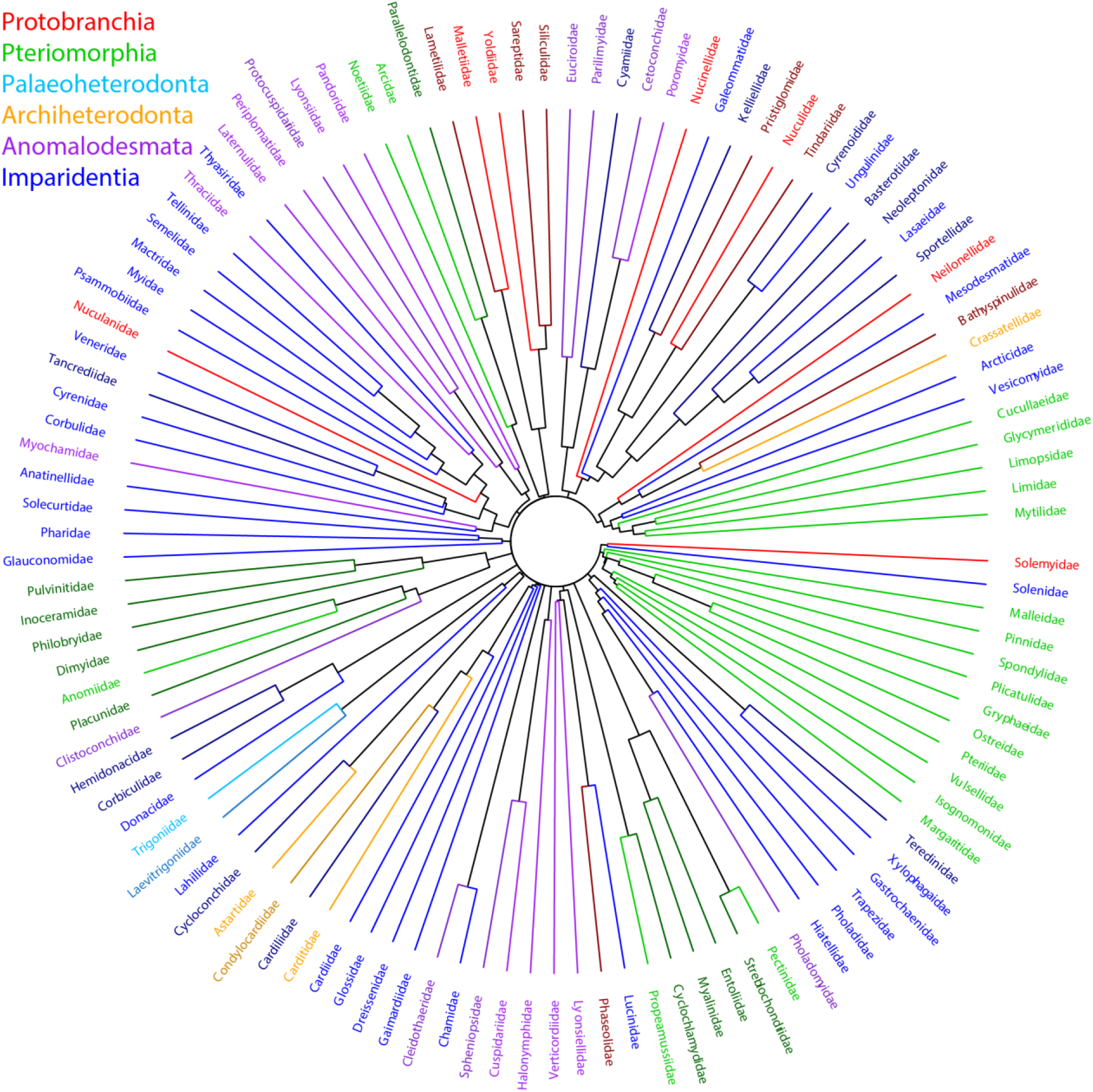
Cluster diagram of the predictions of the family CNN for 74,820 images belonging to 114 families. Color coding by subclass, darker shades indicate target families (those not included in the training).

For the training families, the ensemble approach (Figure 9) always improved their node distances to their respective orders and subclasses, and also the tree coherency index. This does not apply, however, to the node distances of the target families to the orders and subclasses to which they belong according to WoRMS. Combining the ensemble tree with five published phylogenetic trees of the Bivalvia into a supertree (Figure 10) improved the tree coherency index and the overall tree topology. This was expected because the subclass-level topology of the bivalve tree of life is relatively well established and is very similar among the five published trees, and the supertree approach superimposed this topology onto the ensemble tree. The supertree approach also improved the node distances of the target families.

**Figure 9.**
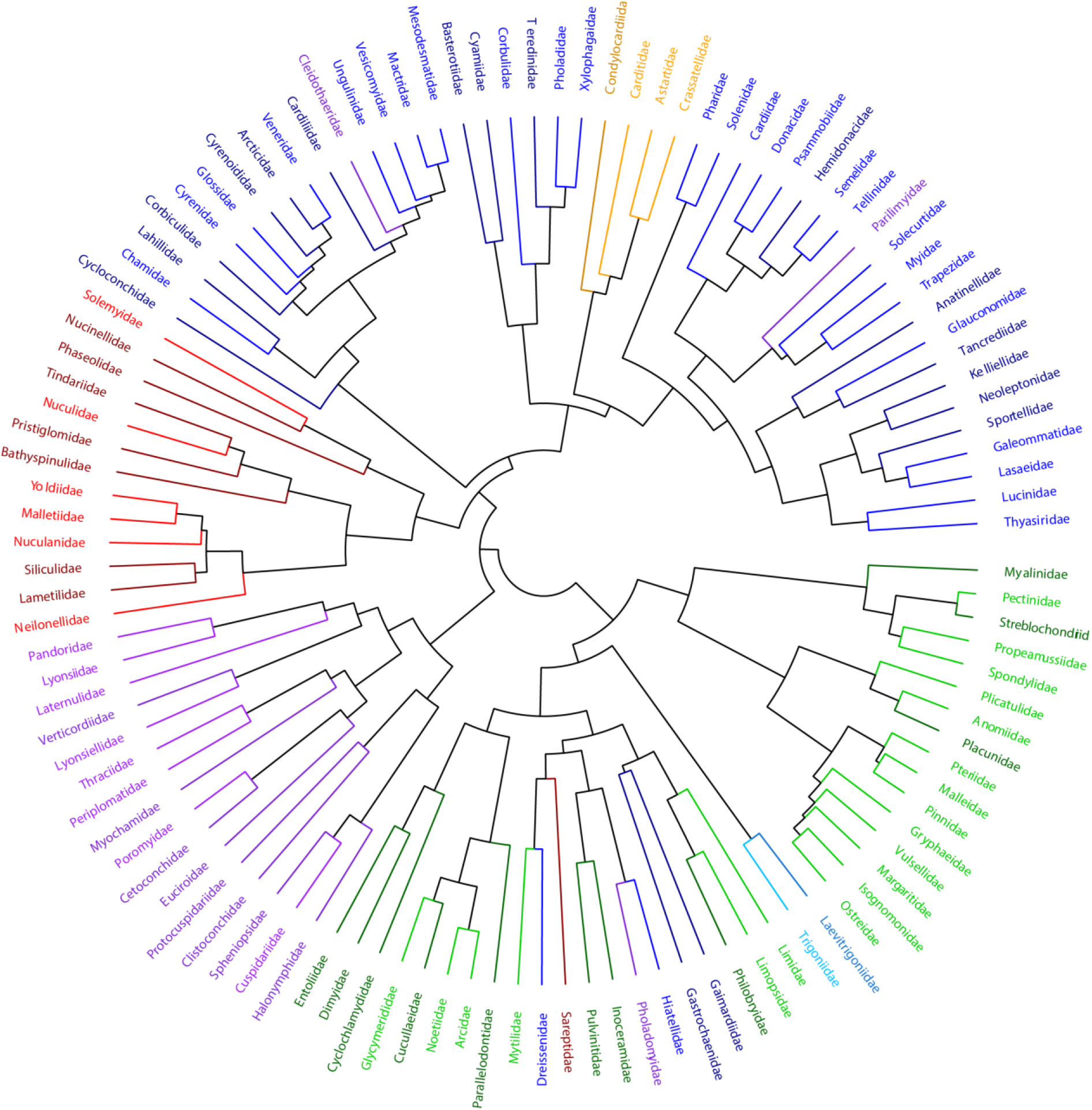
The ensemble tree made from combined predictions of the family, order, and subclass CNNs, each for the same 74,820 images belonging to 114 families. Color coding as in figure 8.

**Figure 10.**
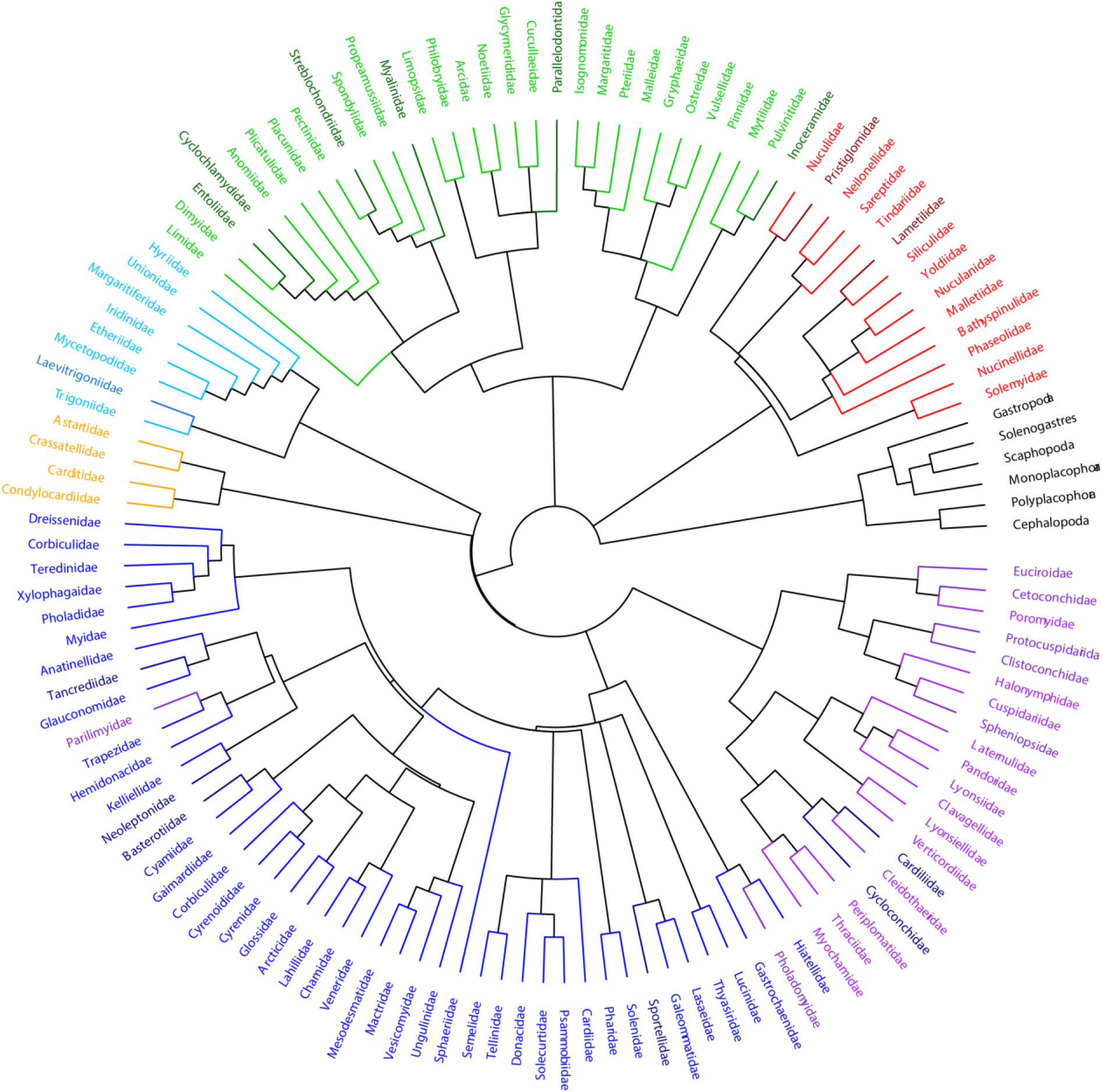
A supertree for 128 bivalve families, combined from the ensemble tree shown in figure 9 and the five published phylogenies listed in Table S1. Color coding as in figure 8.

### Phylogenetic positions of the target families

Twenty-one families of the supertree were neither in the training set nor included in any of the five published phylogenies used for the supertree. Their positions in the supertree are here discussed in more detail.

*Lametilidae (Protobranchia)* is generally placed in the order Nuculanida (Allen & Sanders 1973; Carter et al. 2011). The name-giving species *Lametila abyssorum* was considered as belonging to Phaseolidae (Sharma et al. 2013), but this was questioned by Huber (2015) who retained it as valid family. Here Lametilidae comes out as sister to Siliculidae among other Nuculanida, including Phaseolidae.

*Pristiglomidae (Protobranchia)* come out as sister to Nuculidae, an affinity already proposed when the family was introduced (Sanders & Allen 1973). In the molecular phylogeny by Sharma et al. (2013), three undetermined species of *Pristigloma* fell on a long branch basal to all other Nuculanida and were included in Sareptidae as a separate superfamily Sareptoidea, sister to all other Nuculanoidea. Here, Sareptidae is on a separate branch among the Protobranchia, with Neilonellidae and Tindariidae.

*Cyclochlamydidae (Pteriomorphia)* are micro-scallops recently separated from Propeamussiidae and considered part of Pectinoidea (Dijkstra & Maestrati 2012); molecular data are so far not available (Smedley et al. 2019). Here it is sister to Dimyidae-Entoliidae among the Anomioidea and thus on a separate branch as Pectinidae-Propeamussiidae. Notably, in the tree derived from predictions of the family-CNN alone, Cyclochlamydidae are sister to Propeamussiidae.

*Entoliidae (Pteriomorphia)* is largely an extinct family and superfamily with only two extant species (*Pectinella aequoris* and *P. sigbeei*) and is placed here as sister to Dimyidae. Waller (2006) placed Entoliidae as sister to Propeamussiidae and both as sister to Pectinidae-Spondylidae, whereas Smedley et al. (2019) also retrieved Entoliidae as sister to Propeamussiidae, but both as sister to Spondylidae and the three as sister to Pectinidae. Neither of the two analyses, however, included Dimyidae or any Anomioidea. Close relations of Dimyidae and Plicatulidae, as found here, were rejected based on conchological and shell microstructual features (Hautmann 2001).

*Inoceramidae (Pteriomorphia)* is an enigmatic extinct family that includes some of the largest bivalves that ever lived (Kauffman et al. 2007). Its phylogenetic position within the Bivalvia, as well as its origins, composition, and stratigraphic range, remain controversial (Crame 1982; Crampton 1996; Kauffman & Runnegar 1975; Knight & Morris 2009). It was placed among the extinct superfamilies Praecardioida or Ambonychioidea, or in its own superfamily within Myalinida, and is overall considered to be part of Ostreida (Carter et al. 2011; Knight & Morris 2019; Malchus 2004). Here it appears as sister to Pulvinitidae, on a branch that also includes Mytilidae and is sister to a large branch including Pinnidae, Pteriidae and Ostreidae.

*Myalinidae (Pteriomorphia)* are here found basal among Pectinoidea, which is a quite different position as previously suggested, such as relations to Mytiloidea (Newell 1942; Nevesskaja 2009) or as family within Ambonychoidea, either among Pterioida (Bouchet & Rocroi 2010; McRoberts & Newell 2005), Arcida (Anelli et al. 2006), or the ‘cohort’ Ostreomorphi (Carter et al. 2011).

*Parallelodontidae (Pteriomorphia)* is in all three trees (family-only, ensemble, and supertree) retrieved either basal or sister to other Arcoidea. This supports the view that Glycymerididae and Cucullaeidae may have evolved from a parallelodontid ancestor, as suggested by homologies in ligament and shell microstructural features (Carter 1990; Malchus & Warén 2005; Nicol 1950). Parallelodontidae has a rich Mesozoic fossil record and is either regarded extinct (Bieler et al. 2014; Oliver & Holmes 2006), while others consider *Kamenevus dalli* from Chile as the single surviving parallelodontiid (Newell 1969; Valentich-Scott et al. 2020).

*Streblochondriidae (Pteriomorphia)* is a Mesozoic family commonly placed among the extinct superfamilies Aviculopectinoidea (Amler 1994; Carter & Hautmann 2011; Pagani 2005) or Chaenocardioidea (Carter et al. 2011). It is here retrieved as sister to Pectinidae, supporting the suggested similarities of Streblochondridae with modern pectinids rather than their Paleozoic counterparts (Newell & Boyd 1985) and that Pectinidae may be derived from the Paleozoic-Mesozoic Streblochondriidae (Hautmann 2020). However, Streblochondriidae lack a ctenolium, which is considered a fundamental derived character of modern pectinoids (Waller 1984).

*Laevitrigoniidae* (Palaeoheterodonta) appear as sister to Trigoniidae in all three CNN-derived trees, in agreement with the classification of this family among Trigoniida (Carter et al. 2011).

*Cetoconchidae (Anomalodesmata)* comes out as sister to Poromyidae in all three CNN-derived trees, a relationship supported by shell and morphological characters (Bieler et al. 2010; Dall 1886; Machado 2018; Machado et al. 2019).

*Clistoconchidae (Anomalodesmata)* plot as sister to Protocuspidariidae, which is not supported by anatomical characters indicating thracioid affinities instead (Machado 2018; Morton 2012).

*Euciroidae (Anomalodesmata)* is retrieved as sister to Cetomyidae-Poromyidae, broadly in agreement with the cladistics analyses by Harper et al. (2006), though their molecular data alone suggest relationships to Thraciidae and Myochamidae (Harper et al. 2006, their figs. 4 and 5), which are not supported here. Also not supported here is the traditional classification among Verticordioidea (Newell 1965; Poutiers & Bernard 1995).

*Pholadomyidae and Parilimyidae (Anomalodesmata)* appear in widely different places outside the Anomalodesmata, among Imparidentia families with elongate-quadrate shells (Pholadomyidae with Trapeziidae, Parilimyidae with Hiatellidae). This is frustrating because both families are regarded as stem groups for all extant anomalodesmatans, with a fossil record ranging back to the late Paleozoic (Harper et al. 2006; Harper et al. 2000; Runnegar 1974).

*Protocuspidariidae and Spheniopsidae (Anomalodesmata)* are both currently included in the superfamily Cuspidarioidea (Krylova 1995; Machado et al. 2019). Whereas Spheniopsidae is here retrieved as sister to Cuspidariidae, Protocuspidariidae came out as sister to Clistoconchidae, and both as sister to the superfamily Cuspidarioidea (Halonymphidae, Cuspidariidae, Spheniopsidae).

*Basterotiidae (Imparidentia)* is retrieved as sister to Cyamiidae, where the family has traditionally been placed based on morphological characters (Bieler et al. 2010; Coan & Valentich-Scott 2012). This placement, however, seems in conflict with molecular data indicating affinities of Basterotiidae to Galeommatidae (Giribet & Distel 2003; Goto et al. 2011; Goto et al. 2012; Taylor et al. 2007; Zelaya et al. 2020).

*Cardiliidae (Imparidentia)* plot inconsistently in disparate places in the three CNN-derived trees: in the family-CNN tree it is found among the archiheterodonts Condylocardiidae and Carditidae, and in the ensemble tree it is basal to a branch of mostly Imparidentia including Mactridae and Mesodesmatidae [the traditional place of Cardiliidae (Bieler et al. 2010; Carter et al. 2011)], but also with the anomalodesmatan Cleidothaeidae. In the supertree Cardiliidae is sister to Cleidothaeidae, on a heterogeneous branch within the Anomalodesmata, which also includes the Imparidentia family Cycloconchidae. Although Signorelli & Raven (2018) followed the traditional placement of Cardiliidae among Mactroidea, they considered it uncertain due to recent rearrangements of Mactroidea based on molecular studies (Combosch et al. 2017; Taylor et al. 2007). Unfortunately, the present analysis provides little to improve this situation.

*Neoleptonidae (Imparidentia)* appear as sister to Kelliellidae in both the ensemble and the supertree. Neoleptonidae is commonly placed among Cyamioidea (Morton 2015) but close affinities to Cyamiidae are not seen here. It is notable, though, that in the family-CNN tree it plots close to Sportellidae, a family considered potentially related to Neoleptonidae (Coan 1999; Morton 2015). Close affinities to Veneroidea, as proposed by Salas & Gofas (1998), were not found in any of the CNN-derived trees. A sister-taxa relationship to Kelliellidae, as seen here, is also dubious, as Kelliellidae show close morphological and molecular relationships to Vesicomyidae and Veneridae (Allen 2001; Bieler et al. 2014; Combosch et al. 2017; Krylova et al. 2018).

*Sportellidae (Imparidentia)* are sister to the pair Galeommatidae-Lasaeidae in both ensemble tree and supertree, consistent with the molecular phylogenetic position of this family (Giribet & Distel 2003; Goto et al. 2011; Goto et al. 2012; Taylor et al. 2007). The traditional placement of Sportellidae among Cyamioidea (Coan 1999; Morton 2015; Newell 1965; Thiele 1934) is not supported.

*Tancrediidae (Imparidentia)* is an exclusively fossil family placed either among Tellinoidea (Carter et al. 2011; Chavan 1950; Saul 1989) or Cardioidea (Saul & Popenoe 1962; Speden 1970). Here in both ensemble and supertree, it appears as sister to Glauconomidae, with both being sister to Anatinellidae. This clustering is somewhat unusual, as molecular and morphological data place Glauconomidae among Cyrenoidea (Healy et al. 2006; Mikkelsen et al. 2006; Taylor et al. 2009), whereas Anatinellidae is classified as Mactroidea based on morphological data (Signorelli & Carter 2016); both Cyrenoidea and Mactroidea are only distantly related to Tellinoidea and Cardioidea among the Imparidentia (Bieler et al. 2014; Combosch et al. 2017; Lemer et al. 2019).

## ISSUES, REFLECTIONS, AND FUTURE DIRECTIONS

### Inside-outside views

In conventional shell-based bivalve taxonomy, the outer side of the shell has fewer and less informative characters compared to the inner side, where hinge dentition, muscle scars, pallial line and sinus, etc., are found. It is thus perhaps not surprising that several of the target families for which only images of the outer side of the shell were available, appeared in particularly unsatisfactory positions on the trees (i.e., Inoceramidae, Pholadomyidae). The outer side of many extant bivalve shells, however, is typically more colorful and bears visually attractive sculptural features such as ribs and spines. Hence the websites of collectors and shell dealers, from which many of the images used here have been taken, show the outer side of the shell but not the inside. Indeed, an estimated two thirds of all images used here show the outer side of the shell.

This bias implies that the CNNs trained to a significant extent on the less informative side of the shell. This might have resulted not only in the overall low accuracy of the predictions, reaching 64-77% compared to accuracies beyond 98% for some groups of plants and insects (i.e., Valan et al. 2019). It might also have subdued the ability of the CNNs to recognize similarities among bivalves from highly informative characters such as the hinge dentition, even if the images show them. Besides the obvious way to address this issue –training on more images showing the inside – future approaches might use two or more CNNs in parallel, each trained exclusively on images showing one view of the shell only.

### Systematic differences between prediction distributions

Visual inspection of the family-only and the ensemble tree suggests that target families cluster with other target families more often than one would perhaps expect. Furthermore, the ensemble tree contains a suspicious cluster (Figure 9, center of lower half of tree) composed of numerous target families of mixed taxonomic affinities, including Philobryidae, Gaimardiidae, Inoceramidae, Pulvinitidae, Sareptidae, which gives the impression of being a collection of difficult-to-place ‘leftovers’. Indeed, in the family-only and the ensemble tree, target families more commonly come out as ‘sister taxa’ than expected by chance. This was tested by comparing the actual number of target family pairs (family-only tree: n=7, ensemble tree: n=6) to the number of such pairs in 1000 randomized trees (mean=4.37) using a one-sample t-test; the null hypothesis of equal means was rejected with a confidence of p<0.0001 for both the family-only and the ensemble tree.

The likely underlying reason is a systematic difference in the distribution of the predictions between train and target families. Among their prediction values, each training family has one value that is markedly higher than all others: that for itself. The target families are less likely to have one such distinctive value. Indeed, the highest prediction values among the training families (mean=0.79) are much higher than those of the target families (mean=0.39), which is significantly different from the null hypothesis of equal means (Welch two-sample t-test, p<0.0001). Likewise, the distribution of predictions of the training families is significantly more uneven compared to the target families. Here the Shannon-Wiener index of evenness was calculated for the predictions set of each family among the training and target families, and their medians compared using a Wilcoxon rank sum test (median evenness of training family predictions=0.97, of target family predictions=1.78, which is significantly different, p<0.0001). One potential way to avoid this issue is including the target families in the training process, and to mediate the low number of available images and resulting highly imbalanced training data by employing more sophisticated data augmentation methods (Buda et al. 2018). The downside are the high computational costs. For each additional set of families, the CNNs have to train again on the complete image set, even if the added families comprise only few images.

### Training taxa selection

The rationale for the ensemble approach to address the issue of convergent evolution was the hope that a CNN trained on the morphological diversity of a higher taxonomic unit (i.e., an order or subclass) would be able to identify also families of this higher taxonomic unit that had not been included in the training. In this context it should be pointed out that families were here included in the training set solely based on the number of images available, but neither on how firmly established their phylogenetic positions in the bivalve tree of life are, nor how well these families capture the full morphologic diversity of their respective higher taxa. For example, a CNN trained on Pectinidae and Propeamussiidae is likely to have problems recognizing Anomiidae as member of Pectinida, while a CNN trained on Pectinidae and Anomiidae would most likely recognize Propeamussiidae as member of Pectinida. Considering these points might help to improve this approach in the future.

### Image numbers and node distances

Very low numbers of images could potentially result in an unreliable placement of a family on the tree, because they bear the risk that by pure chance the specimens on the images have strong resembles to members of an entirely unrelated family. While unavoidable in theory, plotting image numbers against node distances of the target families in both ensemble tree and supertree (Figure 11) indicates that this is not a major issue. This observation is promising because a large part of molluscan biodiversity is represented by rare taxa (Bouchet et al. 2002) and several rare and hence little-studied bivalve families are known from only few species or even specimens [i.e., Clistoconchidae (Morton 2012), Galatheavalvidae (Knudsen 1970)].

**Figure 11.**
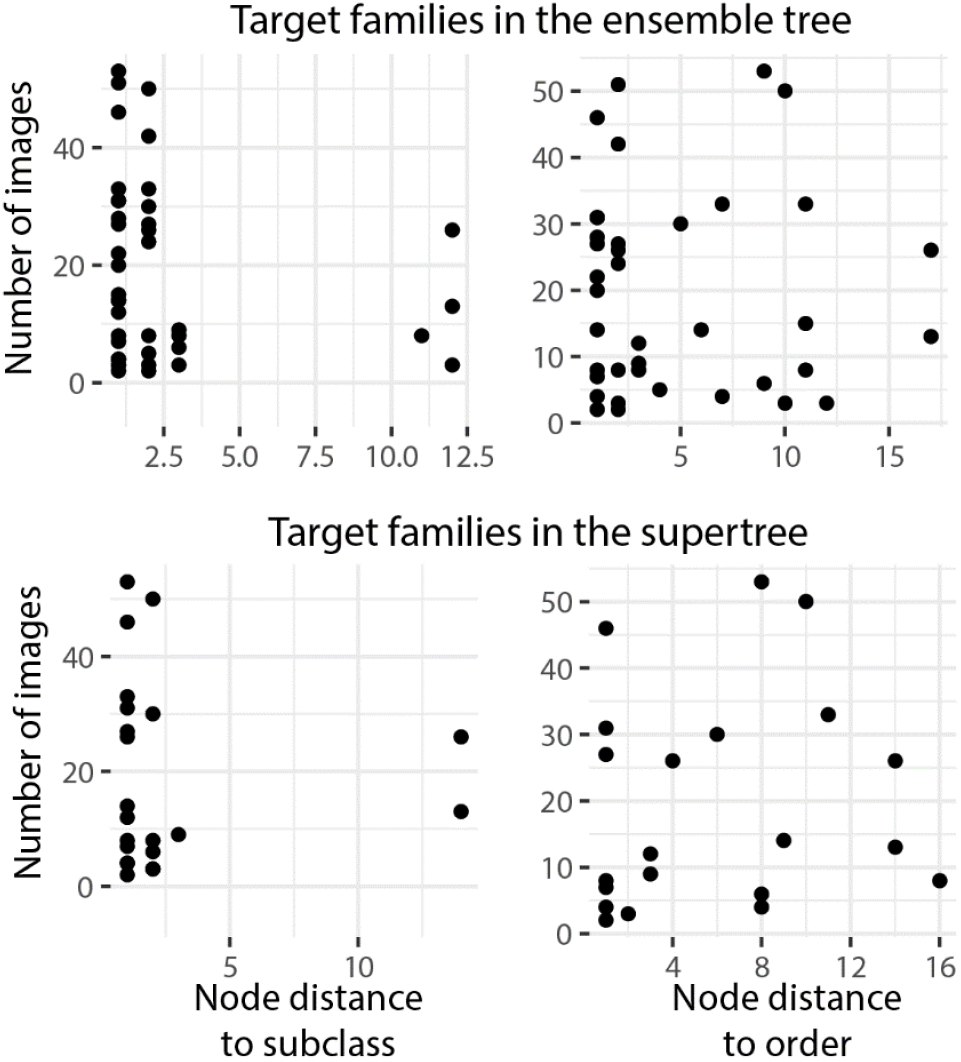
Relationships between node distances and the number of images available for prediction, for the target families in the ensemble and supertree.

### Does increased prediction accuracy improve the inferred phylogenetic relationships?

Put somewhat simplified, the present approach utilizes the identification errors of the CNNs to infer phylogenetic relationships. As pointed out above, if applied to “perfect” predictions, the resulting tree would be a rake without any subgroups. One might thus argue that improved prediction accuracies of the CNNs would not necessarily improve the inferred phylogenetic relationships. However, the improving node distances of the target families through training in the family- and subclass-CNNs indicate that putting more effort into improving the prediction accuracies will also improve the inferred phylogenetic relationships.

## CONCLUSIONS

Deep Learning and Computer Vision methods typically used in image classification, were here explored for their applicability to infer phylogenetic relationships of bivalves. The output (predictions) of convolutional neural networks trained on thousands of images of 75 bivalve families was used to detect and visualize relationships among bivalves based on similarities in the morphology shown on the images. The rather simple approach outlined here is promising, in particular because increasing prediction accuracies of the CNNs appear to result in a better placement of target families for which only few images are available. Thus, methods from Deep Learning and Computer Vision provide a new venue for phylogenetic assessments of large numbers of taxa without labor-intensive hand coding of characters, which is especially relevant for fossil groups. Furthermore, the approach is scalable: it would require only a few modifications to run the same analysis on the 1113 genera to which the species on the images are presently assigned. This scale would hardly be possible when characters had to be hand-coded.

## ACKNOWLEDGMENTS

I would like to thank Miroslav Valan (Stockholm) for inspiration, Johan Nylander (Stockholm) for help with analyzing phylogenetic data, Frans Slieker (Rotterdam) for help with access to the Rotterdam Natural History Museum specimen database, and the WoRMS and MolluscaBase teams for easy access to their data. Financial support was provided by the Swedish Science Foundation (Vetenskapsrådet) through grant 2016-03920.

## REFERENCES

Abadi M, Agarwal A, Barham P, Brevdo E, Chen Z, Citro C, Corrado GS, Davis A, Dean J, Devin M, Ghemawat S, Goodfellow I, Harp A, Irving G, Isard M, Jia Y, Jozefowicz R, Kaiser L, Kudlur M, Levenberg J, Mane D, Monga R, Moore S, Murray D, Olah C, Schuster M, Shlens J, Steiner B, Sutskever I, Talwar K, Tucker P, Vanhoucke V, Vasudevan V, Viegas F, Vinyals O, Warden P, Wattenberg M, Wicke M, Yu Y, and Zheng X. 2016. TensorFlow: large-scale machine learning on heterogeneous distributed systems. Available from arXiv:1603.04467

Allen JA. 2001. The family Kelliellidae (Bivalvia: Heterodonta) from the deep Atlantic and its relationship with the family Vesicomyidae. Zoological Journal of the Linnean Society 131:199–226.

Allen JA, and Sanders HL. 1973. Studies on deep-sea Protobranchia (Bivalvia); the families Siliculidae and Larnetilidae. Bulletin of the Museum of Comparative Zoology, Harvard Collection 145:263–310.

Amler MRW. 1994. The earliest European Streblochondriid bivalves (Pteriomorphia; late Famennian). Annales de la Société géologique de Belgique 117:1–17.

Anelli LE, Rocha-Campos AC, and Simoes MG. 2006. Pennsylvannian pteriomorphian bivalves from the Piauí Formation, Parnaíba Basin, Brazil. Journal of Paleontology 80:1135–1141.

Azizpour H, Razavian AS, Sullivan J, Maki A, and Carlsson S. 2016. Factors of transferability for a generic ConvNet representation. IEEE Transactions on Pattern Analysis and Machine Intelligence 38:1790–1802.

Baraboshkin EE, Ismailova LS, Orlov DM, Zhukovskaya EA, Kalmykov GA, Khotylev OV, Baraboshkin EY, and Koroteev DA. 2020. Deep convolutions for in-depth automated rock typing. Computer & Geosciences 135:104330.

Barré P, Stöver BC, Müller KF, and Steinhage V. 2017. LeafNet: A computer vision system for automatic plant species identification. Ecological Informatics 40:50–56.

Barthelmé S, and Tschumperlé D. 2019. imager: an R package for image processing based on CImg. Journal of Open Source Software 4:1012. https://doi.org/10.21105/joss.01012

Bengio Y. 2011. Deep learning of representations for unsupervised and transfer learning. Proceedings of ICML Workshop on Unsupervised and Transfer Learning Bellevue: IEEE:17–36.

Bieler R, Carter JG, and Coan EV. 2010. Classification of Bivalve families. In: Bouchet P, Rocroi JP, editors. Nomenclator of Bivalve Families. Malacologia 52:113–133.

Bieler R, Mikkelsen PM, Collins TM, Glover EA, Gonzáles VL, Graf DL, Harper EM, Healy JM, Kawauchi GY, Sharma PP, Staubach S, Strong EE, Taylor JD, Temkin I, Zardus JD, Clark S, Guzmán A, McIntyre E, Sharp P, and Giribet G. 2014. Investigating the Bivalve Tree of Life - an exemplar-based approach combining molecular and novel morphological characters. Invertebrate Systematics 28:32–115.

Bouchet P, Lozouet P, Maestrati P, and Heros V. 2002. Assessing the magnitude of species richness in tropical marine environments: exceptionally high numbers of molluscs at a New Caledonia site. Biological Journal of the Linnean Society 75:421–436.

Bouchet P, and Rocroi J-P. 2010. Nomenclator of bivalve families; with a classification of bivalve families by R. Bieler, J.G. Carter & E.V. Coan. Malacologia 52:1–184.

Buda M, Maki A, and Mazurowski MA. 2018. A systematic study of the class imbalance problem in convolutional neural networks. Neural Networks 106:249–259.

Carter JG. 1990. Skeletal biomineralization: patterns, processes and evolutionary trends volume I. New York: Van Nostrand Reinhold.

Carter JG, Altaba CR, Anderson LC, Araujo R, Alexander S. Biakov, Bogan AE, Campbell DC, Campbell M, Jin-hua C, Cope JCW, Delvene G, Dijkstra HH, Zong-jie F, Gardner RN, Gavrilova VA, Irina A. Goncharova, Harries PJ, Hartman JH, Hautmann M, Hoeh WR, Hylleberg J, Bao-yu J, Johnston P, Kirkendale L, Kleemann K, Koppka J, Kříž J, Deusana Machado, Malchus N, Márquez-Aliaga A, Masse J-P, McRoberts CA, Middelfart PU, Mitchell S, Nevesskaja LA, Özer S, Jr. JP, Polubotko IV, Pons JM, Popov S, Sánchez T, Sartori AF, Scott RW, Sey II, Signorelli JH, Silantiev VV, Skelton PW, Steuber T, Waterhouse JB, Wingard GL, and Yancey T. 2011. A synoptical classification of the Bivalvia (Mollusca). Paleontological Contributions 4:1–47.

Carter JG, and Hautmann M. 2011. Shell microstructure of the basal pectinid *Pleuronectites laevigatus*: implications for pectinoid phylogeny (Mollusca: Bivalvia: Pteriomorphia). Journal of Paleontology 85:464–467.

Chavan A. 1950. Remarques sur les Tellinacea du Jurassique superieur. Bulletin de l’Institut royal des Sciences naturelles de Belgique 26:1–19.

Chollet F. 2015. Keras. GitHub. https://github.com/fchollet/keras.

Chollet F. 2016. Xception: deep learning with depthwise separable convolutions. arXiv:1610.02357.

Coan EV. 1999. The eastern Pacific Sportellidae (Bivalvia). The Veliger 42:132–151.

Coan EV, and Valentich-Scott P. 2012. Bivalve Seashells of Tropical West America. Marine Bivalve Mollusks from Baja California to Northern Peru. Santa Barbara: Santa Barbara Museum of Natural History.

Combosch DJ, Collins TM, Glover EA, Graf DL, Harper EM, Healy JM, Kawauchi GY, Lemer S, McIntyre E, Strong EE, Taylor JD, Zardus JD, Mikkelsen PM, Giribet G, and Bieler R. 2017. A family-level Tree of Life for bivalves based on a Sanger-sequencing approach. Molecular Phylogenetics and Evolution 107:191–208.

Crame JA. 1982. Late Jurassic inoceramid bivalves from the Antarctic Peninsula and their stratigraphic use. Palaeontology 25:555–603.

Crame JA. 2002. Evolution of taxonomic diversity gradients in the marine realm: a comparison of Late Jurassic and Recent bivalve faunas. Paleobiology 28:184–207.

Crampton JS. 1996. Inoceramid bivalves from the late Cretaceous of New Zealand. Institute of Geological & Nuclear Sciences Monograph 14:1–192.

Dall WH. 1886. Reports on the results of dredging, under the supervision of Alexander Agassiz, in the Gulf of Mexico (1877-78) and in the Caribbean Sea (1879-80), by the U.S. Coast Survey steamer “Blake” XXIX. Report on the Mollusca. Part 1, Brachiopoda and Pelecypoda. Bulletin of the Museum of Comparative Zoology, Harvard University 12:171–318.

de Lima RP, Duarte D, Nicholson C, Slatt R, and Marfurt KJ. 2020a. Petrographic microfacies classification with deep convolutional neural networks. Computer & Geosciences 104481.

de Lima RP, Suriamin F, Marfurt KJ, and Pranter MJ. 2019. Convolutional neural networks as aid in core lithofacies classification. Interpretation 7:SF27–SF40.

de Lima RP, Welch K, Barrick JE, Marfurt KJ, Burkhalter R, Cassel M, and Soreghan GS. 2020b. Convolutional neural networks as an aid to biostratigraphy and micropaleontology: a test on late Paleozoic microfossils. Palaios 35:391–402.

Deng J, Dong W, Socher R, Li LJ, Li K, and Fei-Fei L. 2009. Imagenet: a large-scale hierarchical image database. Proceedings of the IEEE Conference on Computer Vision and Pattern Recognition:248–255.

Dijkstra HH, and Maestrati P. 2012. Pectinoidea (Mollusca, Bivalvia, Propeamussiidae, Cyclochlamydidae n. fam., Entoliidae and Pectinidae) from the Vanuatu Archipelago. Zoosystema 34:389–408.

Dunn CW, Hejnol A, Matus DQ, Pang K, Browne WE, Smith SA, Seaver E, Rouse GW, Obst M, Edgecombe GD, Sørensen MV, Haddock SHD, Schmidt-Rhaesa A, Okusu A, Kristensen RM, Wheeler WC, Martindale MQ, and Giribet G. 2008. Broad phylogenomic sampling improves resolution of the animal tree of life. Nature 452:745–749.

Erwin DH. 2008. Extinction as the loss of evolutionary history. Proceedings of the National Academy of Sciences of the USA 105, suppl. 1:11520–11527.

Foote M, Crampton JS, Beu AG, Marshall BA, Cooper RA, Maxwell PA, and Matcham I. 2007. Rise and fall of species occupancy in Cenozoic fossil mollusks. Science 318:1131–1134.

Galili T. 2015. dendextend: an R package for visualizing, adjusting and comparing trees of hierarchical clustering. Bioinformatics 31:3718–3720. 10.1093/bioinformatics/btv428

Geng Z, Wu X, Shi Y, and Fomel S. 2020. Deep learning for relative geologic time and seismic horizons. Geophysics 85 87–100.

Giribet G, and Distel DL. 2003. Bivalve phylogeny and molecular data. In: Lydeard C, and Lindberg DR, eds. Molecular systematics and phylogeography of mollusks. Washington: Smithsonian Institution Press, 45–90.

Gonzáles VL, Andrade SCS, Bieler R, Collins TM, Dunn CW, Mikkelsen PM, Taylor JD, and Giribet G. 2015. A phylogenetic backbone for Bivalvia: an RNA-seq approach. Proceedings of the Royal Society B 282:20142332.

Goto R, Hamamura Y, and Kato M. 2011. Morphological and ecological adaptation of *Basterotia* bivalves (Galeommatoidea: Sportellidae) to symbiotic association with burrowing echiuran worms. Zoological Science 28:225–234.

Goto R, Kawakita A, Ishikawa H, Hamamura Y, and Kato M. 2012. Molecular phylogeny of the bivalve superfamily Galeommatoidea (Heterodonta, Veneroida) reveals dynamic evolution of symbiotic lifestyle and interphylum host switching. BMC Evolutionary Biology 12:172.

Gould SJ, and Calloway CB. 1980. Clams and brachiopods-ships that pass in the night. Paleobiology 6:383–396.

Harper EM, Dreyer H, and Steiner G. 2006. Reconstructing the Anomalodesmata (Mollusca: Bivalvia): morphology and molecules. Zoological Journal of the Linnean Society 148:395–420.

Harper EM, Hide EA, and Morton B. 2000. Relationships between the extant Anomalodesmata: a cladistic test. Geological Society, London, Special Publications 177:129–143. https://doi.org/10.1144/GSL.SP.2000.177.01.07

Hautmann M. 2001. Taxonomy and phylogeny of cementing Triassic Bivalves (Families Prospondylidae, Plicatulidae, Dimyidae and Ostreidae). Palaeontology 44:339–373.

Hautmann M. 2020. The first scallop. Paläontologische Zeitschrift 84:317–322.

He K, Zhang X, Ren S, and Sun J. 2015. Deep residual learning for image recognition. https://arxiv.org/abs/1512.03385

Healy JM, Mikkelsen PM, and Bieler R. 2006. Sperm ultrastructure in *Glauconome plankta* and its relevance to the affinities of the Glauconomidae (Bivalvia: Heterodonta). Invertebrate Reproduction & Development 49:29–39. https://doi.org/10.1080/07924259.2006.9652191

Hsiang AY, Brombacher A, Rillo MC, Mleneck‐Vautravers MJ, Conn S, Lordsmith S, Jentzen A, Henehan MJ, Metcalfe B, Fenton IS, Wade BS, Fox L, Meilland J, Davis CV, Baranowski U, Groeneveld J, Edgar KM, Movellan A, Aze T, Dowsett HJ, Miller CG, Rios N, and Hull PM. 2019. Endless Forams: >34,000 modern planktonic foraminiferal images for taxonomic training and automated species recognition using convolutional neural networks. Paleoceanography and Paleoclimatology 34:1157–1177.

Huang J, Rathod V, Sun C, Zhu M, Korattikara A, Fathi A, Fischer I, Wojna Z, Song Y, Guadarrama S, and Murphy K. 2017a. Speed/accuracy trade-offs for modern convolutional object detectors. Proceedings of the IEEE Conference on Computer Vision and Pattern Recognition:7310–7311.

Huang L, Dong X, and Clee TE. 2017b. A scalable deep learning platform for identifying geologic features from seismic attributes. Leading Edge 36:249–256.

Huber M. 2015. Compendium of Bivalves 2. Harxheim: ConchBooks.

Jablonski D, and Finarelli JA. 2009. Congruence of morphologically-defined genera with molecular phylogenies. Proceedings of the National Academy of Sciences of the USA 106:8262–8266.

Jablonski D, Roy K, and Valentine DL. 2006. Out of the tropics: Evolutionary dynamics of the latitudinal diversity gradient. Science 314:102–106.

Jablonski D, Sepkoski JJ, Bottjer DJ, and Sheehan PM. 1983. Onshore-offshore patterns in the evolution of Phanerozoic shelf communities. Science 222:1123–1125.

John C, and Kanagandran S. 2019. AI to improve the reliability and reproducibility of descriptive data: a case study using convolutional neural networks to recognize carbonate facies in cores. AAPG Annual Convention and Exhibition. San Antonio, TX.

Jombart T, Balloux F, and Dray S. 2010. *adephylo*: new tools for investigating the phylogenetic signal in biological traits. Bioinformatics 26:1907–1909. https://doi.org/10.1093/bioinformatics/btq292

Kauffman EG, Harries PJ, Meyer C, Villamil T, Arango C, and Jaecks G. 2007. Paleoecology of giant Inoceramidae (*Platyceramus*) on a Santonian (Cretaceous) seafloor in Colorado. Journal of Paleontology 81:64–81.

Kauffman EG, and Runnegar BN. 1975. *Atomodesma* (Bivalvia), and Permian species of the United States. Journal of Paleontology 49:23–41.

Knight RI, and Morris NJ. 2009. A reconsideration of the origins of the ‘typical’ Cretaceous inoceramid calcitic hinge plate in the light of new ultrastructural observations from some Jurassic ‘inoceramids’. Palaeontology 52:963–989.

Knight RI, and Morris NJ. 2019. Well-developed muscle attachments in British Albian inoceramids (Inoceramidae, Bivalvia): Implications for inoceramid paleobiology, evolution and taxonomy. Papers in Palaeontology 5:461–481.

Knope ML, Bush AM, Frishkoff LO, Heim NA, and Payne JL. 2020. Ecologically diverse clades dominate the oceans via extinction resistance. Science 367:1035–1038.

Knudsen J. 1970. The systematics and biology of abyssal and hadal Bivalvia. Galathea report 11:1–241.

Koeshidayatullah A, Morsilli M, Lehrmann DJ, Al-Ramadan K, and Payne JL. 2020. Fully automated carbonate petrography using deep convolutional neural networks. Marine and Petroleum Geology 122 104687.

Kroh A, and Smith AB. 2010. The phylogeny and classification of post-Palaeozoic echinoids. Journal of Systematic Palaeontology 8:147–212. 10.1080/14772011003603556

Krylova EM. 1995. Clams of the family Protocuspidariidae (Septibranchia, Cuspidarioidea): taxonomy and distribution. Zoological Journal 74:20–38 (in Russian, with English summary).

Krylova EM, Sahling H, and Borowski C. 2018. Resolving the status of the families Vesicomyidae and Kelliellidae (Bivalvia: Venerida), with notes on their ecology. Journal of Molluscan Studies. 10.1093/mollus/eyx050

Laubach T, Haeseler Av, and Lercher MJ. 2012. TreeSnatcher plus: capturing phylogenetic trees from images. BMC Bioinformatics 13:110.

Laumer CE, Fernández R, Lemer S, Combosch D, Kocot KM, Riesgo A, Andrade SCS, Sterrer W, Sørensen MV, and Giribet G. 2019. Revisiting metazoan phylogeny with genomic sampling of all phyla. Proceedings of the Royal Society B 286:20190831.

LeCun Y, Bengio Y, and Hinton G. 2015. Deep learning. Nature 521:436–444.

Lemer S, Bieler R, and Giribet G. 2019. Resolving the relationships of clams and cockles: dense transcriptome sampling drastically improves the bivalve tree of life. Proceedings of the Royal Society B 286:20182684. http://dx.doi.org/10.1098/rspb.2018.2684

Lockwood R. 2003. Abundance not linked to survival across the Cretaceous mass extinction: Patterns in North American bivalves. Proceedings of the National Academy of Sciences of the USA 100:2478–2482.

Lucas A, and Gautier L. 2020. ctc: Cluster and Tree Conversion. R package version 1.64.0.

Machado FM. 2018. Unravelling the diversity of Anomalodesmata (Mollusca: Bivalvia): a morphological and phylogenetic approach PhD. Universidade Estadual de Campinas.

Machado FM, Passos FD, and Giribet G. 2019. The use of micro-computed tomography as a minimally invasive tool for anatomical study of bivalves (Mollusca: Bivalvia). Zoological Journal of the Linnean Society 186:46–75.

Malchus N. 2004. Early ontogeny of Jurassic bakevelliids and their bearing on bivalve evolution. Acta Geologica Polonica 49:85–110.

Malchus N, and Warén A. 2005. Shell and hinge morphology of juvenile *Limopsis* (Bivalvia: Arcoida) - implications for limopsid evolution. Marine Biology Research 1:350 - 364.

Mata-Montero E, and Carranza-Rojas J. 2016. Automated Plant Species Identification: Challenges and Opportunities. Cham: Springer International Publishing. p 26–36.

McRoberts CA, and Newell ND. 2005. Marine Myalinidae (Bivalvia: Pterioida) from the Permian of West Texas. American Museum Novitates 3469:1–15.

Mikkelsen PM, Bieler R, Kappner I, and Rawlings TA. 2006. Phylogeny of Veneroidea (Mollusca: Bivalvia) based on morphology and molecules. Zoological Journal of the Linnean Society 148:439–521.

Morton B. 2012. The functional morphology and inferred biology of the enigmatic South African “quadrivalve” bivalve *Clistoconcha insignis* Smith, 1910 (Thracioidea: Clistoconchidae fam. nov.): Another anomalodesmatan evolutionary eccentric. Transactions of the Royal Society of South Africa 67:59–89. https://doi.org/10.1080/0035919X.2012.702321

Morton B. 2015. The biology and functional morphology of the placental embryo-brooding *Neolepton salmoneum*, a comparison with *Neolepton subtrigonum* (Bivalvia: Cyamioidea: Neoleptonidae), and a discussion of affinities. American Malacological Bulletin 33:1–21.

Newell ND. 1942. Late Paleozoic pelecypods: Mytilacea. University of Kansas Publications 10:1–80.

Newell ND. 1965. Classification of the Bivalvia. American Museum Novitates 2206:1–25.

Newell ND. 1969. Classification of Bivalvia. In: Moore RC, ed. Treatise on Invertebrate Paleontology, Part N, Mollusca 6 vol 1 Bivalvia. Boulder-Lawrence: Geological Society of America & University of Kansas, N205–N244.

Newell ND, and Boyd DW. 1985. Permian scallops of the pectinacean family Streblochondriidae. American Museum Novitates 2831:1–13.

Nevesskaja LA. 2009. Principles of systematics and the system of bivalves. Paleontological Journal 43:1–11.

Nicol D. 1950. Origin of the pelecypod family Glycymeridae. Journal of Paleontology 24:89–98.

Oksanen J, Kindt R, Legendre P, and O’Hara RB. 2005. vegan: community ecology package.http://cran.r-project.org/.

Oliver PG, and Holmes AM. 2006. The Arcoidea (Mollusca: Bivalvia): a review of the current phenetic-based systematics. Zoological Journal of the Linnean Society 148:237–251.

Ooms J, and Team TID. 2020. Magick: advanced graphics and image-processing in R.https://cran.r-project.org/web/packages/magick/index.html.

Pagani MA. 2005. Los bivalvos carboníferos y pérmicos de la Patagonia (Chubut, Argentina). Parte III: Familias Mytilidae, Pterineidae, Limidae, Leptochondriidae, Etheripectinidae, Euchondriidae y Streblochondriidae. Ameghiniana 42:579–596.

Paradis E, Claude J, and Strimmer K. 2004. APE: Analyses of Phylogenetics and Evolution in R language. Bioinformatics 20:289–290. 10.1093/bioinformatics/btg412

Patzkowsky ME. 2017. Origin and evolution of regional biotas: a deep-time perspective. Annual Review of Earth and Planetary Sciences 45:471–495.

Perez L, and Wang J. 2017. The effectiveness of data augmentation in image classification using deep learning. arXiv:1712.04621.

Peters SE, and Foote M. 2001. Biodiversity in the Phanerozoic: a reinterpretation. Paleobiology 27:583–601.

Pick KS, Philippe H, Schreiber F, Erpenbeck D, Jackson DJ, Wrede P, Wiens M, Alié A, Morgenstern B, Manuel M, and Wörheide G. 2010. Improved phylogenomic taxon sampling noticeably affects nonbilaterian relationships. Molecular Biology and Evolution 27:1983–1987.

Plazzi F, and Passamonti M. 2010. Towards a molecular phylogeny of Mollusks: Bivalves’ early evolution as revealed by mitochondrial genes. Molecular Phylogenetics and Evolution 57:641–657.

Poutiers J-M, and Bernard FR. 1995. Carnivorous bivalve molluscs (Anomalodesmata) from the tropical western Pacific Ocean, with a proposed classification and a catalogue of Recent species. Mémoires de Muséum National de Histoire Naturelle 167:107–187.

Ragan MA. 1992. Phylogenetic inference based on matrix representation of trees. Molecular Phylogenetics and Evolution 1:3–10.

Raup DM, and Jablonski D. 1993. Geography of End-Cretaceous marine bivalve extinctions. Science 260:971–973.

Renema W, Bellwood DR, Braga JC, Bromfield K, Hall R, Johnson KG, Lunt P, Meyer CP, McMonagle LB, Morley RJ, O’Dea A, Todd JA, Wesselingh FP, Wilson MEJ, and Pandolfi JM. 2008. Hopping Hotspots: Global shifts in marine biodiversity. Science 321:654–657.

Runnegar BN. 1974. Evolutionary history of the bivalve subclass Anomalodesmata. Journal of Paleontology 48:904–939.

Sahoo PK, Soltani S, Wong AKC, and Chen YC. 1988. A survey of thresholding techniques. Computer Vision, Graphics, and Image Processing 41:223–260.

Salas C, and Gofas S. 1998. Description of four new species of *Neolepton* Monterosato, 1875 (Mollusca: Bivalvia: Neoleptonidae), with comments on the genus and on its affinity with the Veneracea. Ophelia 48:35–70. http://dx.doi.org/10.1080/00785236.1998.10428676

Sanders HL, and Allen JA. 1973. Studies on deep-sea Protobranchia (Bivalvia) Prologue and the Pristiglomidae. Bulletin of the Museum of Comparative Zoology, Harvard Collection 145:237–262. http://biodiversitylibrary.org/page/4336413

Saul LR. 1989. California Late Cretaceous donaciform bivalves. The Veliger 32:188–208.

Saul LR, and Popenoe WP. 1962. *Meekia*, enigmatic Cretaceous pelecypod genus. University of California Publications in Geological Sciences 40:289–343.

Schliep KP. 2011. phangorn: phylogenetic analysis in R. Bioinformatics 27:592–593

Selvaraju RR, Cogswell M, Das A, Vedantam R, Parikh D, and Batra D. 2017. Grad-CAM: Visual explanations from deep networks via gradient-based localization. Proceedings of the IEEE International Conference on Computer Vision:618–626.

Sepkoski JJ. 1981. A factor analytic description of the Phanerozoic marine fossil record. Paleobiology 7:36–53.

Sharma PP, González VL, Kawauchi GY, Andrade SCS, Guzmán A, Collins TM, Glover EA, Harper EM, Healy JM, Mikkelsen PM, Taylor JD, Bieler R, and Giribet G. 2012. Phylogenetic analysis of four nuclear protein-encoding genes largely corroborates the traditional classification of Bivalvia (Mollusca). Molecular Phylogenetics and Evolution 65:64–74.

Sharma PP, Zardus JD, Boyle EE, González VL, Jennings RM, McIntyre E, Wheeler W, Etter RJ, and Giribet G. 2013. Into the deep: A phylogenetic approach to the bivalve subclass Protobranchia. Molecular Phylogenetics and Evolution 69:188–204.

Signorelli JH, and Carter JG. 2016. The Anatinellidae and Kymatoxinae: A reassessment of their affinities within the superfamily Mactroidea (Mollusca, Bivalvia). American Malacological Bulletin 33:204–211.

Signorelli JH, and Raven H. 2018. Current knowledge of the family Cardiliidae (Bivalvia, Mactroidea). Journal of Paleontology 92:130–145.

Simonyan K, and Zisserman A. 2014. Very deep convolutional networks for large-scale image recognition. arXiv:1409.1556.

Smedley GD, Audino JA, Grula C, Porath-Krause A, Pairett AN, Alejandrino A, Lacey L, Masters F, Duncan PF, Strong EE, and Serb JM. 2019. Molecular phylogeny of the Pectinoidea (Bivalvia) indicates Propeamussiidae to be a non-monophyletic family with one clade sister to the scallops (Pectinidae). Molecular Phylogenetics and Evolution 137:293–299.

Speden IG. 1970. The type Fox Hills Formation, Cretaceous (Maestrichtian), South Dakota. Part 2. Systematics of the Bivalvia. Peabody Museum of Natural History Yale University, Bulletin 33:1–222.

Stanley SM. 1970. Relation of shell form to life habits of the Bivalvia (Mollusca). Geological Society of America Memoir 125:1–282.

Sutskever I, Martens J, Dahl G, and Hinton G. 2013. On the importance of initialization and momentum in deep learning. Proceedings of the 30^th^ International Conference on Machine Learning. Atlanta. p 1139–1147.

Szegedy C, Ioffe S, Vanhoucke V, and Alemi A. 2016. Inceptionv4, Inception-ResNet and the impact of residual connections on learning. arXiv:1602.07261.

Taylor JD, Glover EA, and Williams ST. 2009. Phylogenetic position of the bivalve family Cyrenoididae—removal from (and further dismantling of) the superfamily Lucinoidea. The Nautilus 123:9–13.

Taylor JD, Williams ST, Glover EA, and Dyal P. 2007. A molecular phylogeny of heterodont bivalves (Mollusca: Bivalvia: Heterodonta): new analyses of 18S and 28S rRNA genes. Zoologica Scripta 36:587–606.

Thiele J. 1934. Handbuch der systematischen Weichtierkunde 3. Jena: Gustav Fischer.

Thuy B, and Stöhr S. 2016. A new morphological phylogeny of the Ophiuroidea (Echinodermata) accords with molecular evidence and renders microfossils accessible for cladistics. PLoS ONE 11:e0156140.

Valan M, Mokonyi K, Maki A, Vondráček D, and Ronquist F. 2019. Automated taxonomic identification of insects with expert-level accuracy using effective feature transfer from convolutional networks. Systematic Biology 68:876–895.

Valentich-Scott P, Coan EV, and Zelaya DG. 2020. Bivalve seashells of western South America. Marine bivalve mollusks from Punta Aguja, Perú to Isla Chiloé, Chile. Santa Barbara Museum of Natural History Monographs 8:593.

Valentine JW, Jablonski D, Kidwell SM, and Roy K. 2006. Assessing the fidelity of the fossil record by using marine bivalves. Proceedings of the National Academy of Sciences of the USA 103:6599–6604.

Waller TR. 1984. The ctenolium of scallop shells: functional morphology and evolution of a key family-level character in the Pectinacea (Mollusca: Bivalvia). Malacologia 25:203–219.

Waller TR. 2006. Phylogeny of families in the Pectinoidea (Mollusca: Bivalvia): importance of the fossil records. Zoological Journal of the Linnean Society 148:313–342.

Wäldchen J, and Mäder P. 2018. Machine learning for image based species identification. Methods in Ecology and Evolution. https://doi.org/10.1111/2041-210X.13075

Wäldchen J, Rzanny M, Seeland M, and Mäder P. 2018. Automated plant species identification—Trends and future directions. PLoS Computational Biology 14:e1005993.

Yosinski J, Clune J, Bengio Y, and Lipson H. 2014. How transferable are features in deep neural networks? In: Ghahramani Z, Welling M, Cortes C, Lawrence ND, and Weinberger KQ, eds. Advances in Neural Information Processing Systems: Curran Associates, Inc., 3320–3328.

Zelaya DG, Güller M, and Ituarte C. 2020. Filling a blank in bivalve taxonomy: an integrative analysis of Cyamioidea (Mollusca: Bivalvia). Zoological Journal of the Linnean Society 190:558–591. https://doi.org/10.1093/zoolinnean/zlz144

