## Supplemental Table S1 for "Assessing bivalve phylogeny using Deep Learning and Computer Vision approaches"

**Table S1.** List of families, their classification, image numbers, and usage.

| Subclass | Order | Family | Number of images | Training family | Present in phylogenies for supertree | | | | |
|---|---|---|---|---|---|---|---|---|---|
| | | | | | Bieler | Combosch | Gonzales | Plazzi | Sharma |
| Protobranchia | Nuculanida | Bathyspinulidae | 8 | no | no | yes | no | no | no |
| | | Lametilidae | 8 | no | no | no | no | no | no |
| | | Malletiidae | 182 | yes | yes | yes | no | no | yes |
| | | Neilonellidae | 128 | yes | no | yes | no | no | no |
| | | Nuculanidae | 1239 | yes | yes | yes | no | yes | yes |
| | | Phaseolidae | 5 | no | no | yes | no | no | no |
| | | Pristiglomidae | 26 | no | no | no | no | no | no |
| | | Sareptidae | 8 | no | no | yes | no | no | no |
| | | Siliculidae | 14 | no | no | yes | no | no | no |
| | | Tindariidae | 51 | no | no | yes | no | no | no |
| | | Yoldiidae | 670 | yes | yes | yes | yes | no | yes |
| | Nuculida | Nuculidae | 1195 | yes | yes | yes | yes | yes | yes |
| | Solemyida | Nucinellidae | 62 | yes | yes | yes | no | no | no |
| | | Solemyidae | 161 | yes | yes | no | yes | yes | yes |
| Pteriomorphia | Arcida | Arcidae | 4375 | yes | yes | yes | yes | yes | yes |
| | | Cucullaeidae | 264 | yes | no | no | no | no | no |
| | | Glycymerididae | 2035 | yes | yes | yes | no | no | yes |
| | | Limopsidae | 424 | yes | yes | yes | no | no | no |
| | | Noetiidae | 425 | yes | yes | yes | no | no | yes |
| | | Parallelodontidae | 9 | no | no | no | no | no | no |
| | | Philobryidae | 22 | no | no | yes | yes | no | no |
| | Limida | Limidae | 1497 | yes | yes | yes | no | yes | yes |
| | Myalinida | Inoceramidae | 53 | no | no | no | no | no | no |
| | | Myalinidae | 6 | no | no | no | no | no | no |
| | Mytilida | Mytilidae | 4095 | yes | yes | yes | yes | yes | yes |
| | Ostreida | Gryphaeidae | 428 | yes | yes | yes | no | yes | no |
| | | Isognomonidae | 460 | yes | yes | yes | no | no | no |
| | | Malleidae | 190 | yes | yes | yes | no | no | yes |
| | | Margaritidae | 333 | yes | no | no | yes | yes | no |
| | | Ostreidae | 1636 | yes | yes | yes | no | yes | no |
| | | Pinnidae | 834 | yes | yes | yes | yes | yes | yes |
| | | Pteriidae | 491 | yes | yes | yes | no | no | yes |
| | | Pulvinitidae | 15 | no | yes | yes | no | no | no |
| | | Vulsellidae | 245 | yes | no | no | no | no | no |
| | Pectinida | Anomiidae | 730 | yes | yes | yes | no | yes | yes |
| | | Cyclochlamydidae | 3 | no | no | no | no | no | no |
| | | Dimyidae | 20 | no | yes | yes | no | no | no |
| | | Entoliidae | 7 | no | no | no | no | no | no |
| | | Pectinidae | 8650 | yes | yes | yes | yes | yes | yes |
| | | Placunidae | 31 | no | yes | yes | no | no | no |
| | | Plicatulidae | 407 | yes | yes | yes | no | no | no |
| | | Propeamussiidae | 969 | yes | yes | yes | no | no | no |
| | | Spondylidae | 1863 | yes | yes | yes | no | yes | no |
| | | Streblochondriidae | 2 | no | no | no | no | no | no |
| Paleoheterodonta | Trigoniida | Laevitrigoniidae | 31 | no | no | no | no | no | no |
| | | Trigoniidae | 180 | yes | yes | yes | yes | no | yes |
| Archiheterodonta | Carditida | Astartidae | 1022 | yes | yes | yes | yes | yes | yes |
| | | Carditidae | 1909 | yes | yes | yes | yes | yes | no |
| | | Condylocardiidae | 27 | no | no | yes | no | no | no |
| | | Crassatellidae | 688 | yes | yes | yes | yes | no | yes |
| Anomalodesmata | Cuspidarioidea | Cuspidariidae | 989 | yes | yes | yes | no | yes | yes |
| | | Halonymphidae | 70 | yes | no | no | no | no | no |
| | | Protocuspidariidae | 12 | no | no | no | no | no | no |
| | | Spheniopsidae | 4 | no | no | no | no | no | no |
| | Laternuloidea | Laternulidae | 175 | yes | yes | yes | no | no | no |
| | Myochamoidea | Cleidothaeridae | 3 | no | yes | yes | no | no | no |
| | | Myochamidae | 101 | yes | yes | yes | yes | no | no |
| | Pandoroidea | Lyonsiidae | 174 | yes | yes | yes | yes | no | yes |
| | | Pandoridae | 281 | yes | yes | yes | no | yes | no |
| | Pholadomyoidea | Parilimyidae | 13 | no | no | no | no | no | no |

| | | | | | | | | | |
|---|---|---|---|---|---|---|---|---|---|
| | | Pholadomyidae | 26 | no | no | no | no | no | no |
| | Poromyoidea | Cetoconchidae | 27 | no | no | no | no | no | no |
| | | Poromyidae | 172 | yes | yes | yes | no | no | no |
| | Thracioidea | Clistoconchidae | 4 | no | no | no | no | no | no |
| | | Periplomatidae | 172 | yes | yes | yes | no | no | no |
| | | Thraciidae | 310 | yes | yes | yes | no | yes | yes |
| | Verticordioidea | Euciroidae | 30 | no | no | no | no | no | no |
| | | Lyonsiellidae | 136 | yes | no | no | no | no | no |
| | | Verticordiidae | 84 | yes | yes | yes | no | no | no |
| Imparidentia | Actinodontida | Cycloconchidae | 3 | no | no | no | no | no | no |
| | Adapedonta | Hiatellidae | 431 | yes | yes | yes | yes | yes | no |
| | | Pharidae | 768 | yes | yes | yes | no | yes | no |
| | | Solenidae | 377 | yes | yes | yes | no | no | yes |
| | Cardiida | Cardiidae | 5099 | yes | yes | yes | yes | yes | yes |
| | | Donacidae | 1438 | yes | yes | yes | no | yes | no |
| | | Psammobiidae | 1375 | yes | yes | yes | no | no | no |
| | | Semelidae | 1131 | yes | yes | yes | no | no | no |
| | | Solecurtidae | 327 | yes | yes | yes | no | no | yes |
| | | Tancrediidae | 14 | no | no | no | no | no | no |
| | | Tellinidae | 5270 | yes | yes | yes | yes | no | yes |
| | Cyamioidea | Cyamiidae | 3 | no | yes | yes | no | no | no |
| | | Sportellidae | 50 | no | no | no | no | no | no |
| | Gaimardioidea | Gaimardiidae | 96 | yes | yes | yes | no | no | no |
| | Galeommatida | Basterotiidae | 33 | no | no | no | no | no | no |
| | | Galeommatidae | 180 | yes | yes | yes | yes | no | yes |
| | | Lasaeidae | 906 | yes | yes | yes | yes | no | no |
| | Gastrochaenida | Gastrochaenidae | 68 | yes | yes | yes | yes | no | no |
| | Lucinida | Lucinidae | 3458 | yes | yes | yes | yes | yes | yes |
| | | Thyasiridae | 381 | yes | yes | yes | no | no | yes |
| | Myida | Corbulidae | 1124 | yes | yes | yes | no | no | yes |
| | | Dreissenidae | 133 | yes | yes | yes | no | yes | yes |
| | | Myidae | 347 | yes | yes | yes | yes | yes | yes |
| | | Pholadidae | 645 | yes | yes | yes | no | no | no |
| | | Teredinidae | 42 | no | yes | yes | no | no | yes |
| | | Xylophagaidae | 134 | yes | no | no | no | no | no |
| | Venerida | Anatinellidae | 64 | yes | no | no | no | no | no |
| | | Arcticidae | 143 | yes | yes | yes | yes | no | no |
| | | Cardiliidae | 8 | no | no | no | no | no | no |
| | | Chamidae | 860 | yes | yes | yes | no | no | no |
| | | Corbiculidae | 2 | no | yes | no | yes | no | yes |
| | | Cyrenidae | 363 | yes | no | yes | yes | no | no |
| | | Cyrenoididae | 24 | no | yes | yes | yes | no | yes |
| | | Glauconomidae | 82 | yes | yes | yes | no | no | yes |
| | | Glossidae | 380 | yes | yes | yes | yes | no | yes |
| | | Hemidonacidae | 33 | no | yes | yes | no | no | yes |
| | | Kelliellidae | 28 | no | yes | yes | no | no | no |
| | | Lahillidae | 115 | yes | no | no | no | no | no |
| | | Mactridae | 2260 | yes | yes | yes | no | yes | no |
| | | Mesodesmatidae | 404 | yes | no | yes | no | no | no |
| | | Neoleptonidae | 46 | no | no | no | no | no | no |
| | | Trapezidae | 131 | yes | yes | yes | no | no | no |
| | | Ungulinidae | 444 | yes | yes | yes | yes | no | no |
| | | Veneridae | 4462 | yes | yes | yes | yes | yes | no |
| | | Vesicomyidae | 222 | yes | yes | yes | no | no | no |
